# Exercise induces time-dependent but not sex-specific transcriptomic changes in healthy human skeletal muscle

**DOI:** 10.1101/2025.08.17.667354

**Authors:** Dale F. Taylor, Elizabeth G. Reisman, Andrew Garnham, Romain Barrès, Nolan J. Hoffman, John A. Hawley, Nikeisha J. Caruana, David J. Bishop

## Abstract

Elucidating the time-dependent transcriptional response of skeletal muscle to exercise is essential for uncovering the molecular mechanisms that drive its health-promoting effects. However, previous studies have been limited by a small number of muscle biopsies, often collected at arbitrary time points post-exercise, and in predominantly male subjects. Using the most comprehensive skeletal muscle biopsy time course following a single session of exercise in both males and females, we identified over 16,600 differentially expressed genes (DEGs), including more than 7,000 novel exercise-responsive genes. Of these DEGs, the magnitude of differential expression in 60% of genes was influenced by cardiorespiratory fitness. Although most mitochondrial genes were differentially expressed after exercise, the majority were downregulated at 24–48 hours, suggesting that mitochondrial protein expression may be regulated by post-transcriptional regulation. Despite 1,193 genes showing sex-specific expression at baseline, exercise-induced gene expression differences were minimal between males and females, suggesting that when cardiorespiratory fitness and exercise stimulus are matched, skeletal muscle adaptations are similar between sexes. To enhance data accessibility, we created an interactive Shiny app (https://BishopLab.shinyapps.io/EXERgene/) that allows users to investigate specific genes and interrogate potential mechanisms of skeletal muscle adaptation to exercise.

## Introduction

Exercise is a potent, non-pharmacological intervention that promotes numerous health benefits important for the prevention and treatment of many diseases including type 2 diabetes, cancer, and several neurological disorders^1^. While the health benefits of exercise are well-established, the underlying molecular mechanisms driving these effects remain poorly understood^2^. To facilitate exercise-induced skeletal muscle adaptations, contraction initiates a range of time-dependent molecular responses^3^, which include the activation of transcription factors^4^, the transcription of mRNA from DNA, and translation of mRNA into protein. Although transcription has long been considered the key regulatory step driving functional protein-level adaptations^5,6^, a comprehensive understanding of the exercise-induced transcriptional response remains far from complete due to most studies sampling muscle at only two or three arbitrary time points within the first few hours post-exercise^7–13^. This is a major oversight^14^ and suggests that thousands of exercise-responsive genes may have either been mischaracterised or overlooked entirely in previous studies.

Our understanding of the transcriptional response to exercise is further hindered by the historic underrepresentation of female participants within research^15^. Nonetheless, it has been hypothesised that some of the physiological differences observed between males and females, such as muscle mass, substrate utilisation during exercise, and muscle fibre type ratios^16^, may contribute to sex-specific transcriptomic responses to exercise. Despite several recent transcriptomic studies using RNA-seq reporting sex-specific skeletal muscle gene expression at baseline^17–22^, there is little direct evidence to support the common assumption that biological sex influences exercise-induced gene expression.

A complete temporal map of the exercise-induced transcriptome, which addresses the male bias in the literature, is clearly warranted to better understand the molecular underpinnings of adaptations in skeletal muscle to exercise. To address this knowledge gap, we generated the most comprehensive time course to date of exercise-induced gene expression, with nine biopsies collected from 20 males and 20 females – before (PRE), during (MID), immediately post (+0 h), and +3, +6, +9, +12, +24, and +48 h following a single session of high-intensity interval exercise (HIIE). Collectively, the findings of this study provide new insights into the dynamic temporal responses of exercise-induced gene expression and challenge the accepted dogma that there are meaningful differences in exercise-induced transcription between sexes.

## Results

### An emphasis on early post-exercise time points has prevented the detection of thousands of exercise-induced genes in muscle

To comprehensively investigate exercise-induced gene expression, RNA-seq was performed on skeletal muscle biopsies collected from 20 healthy untrained males and 20 healthy untrained females across nine different time points before, during, and after a single session of high-intensity interval exercise (HIIE) (**Fig. 1a**).

**Fig. 1:**
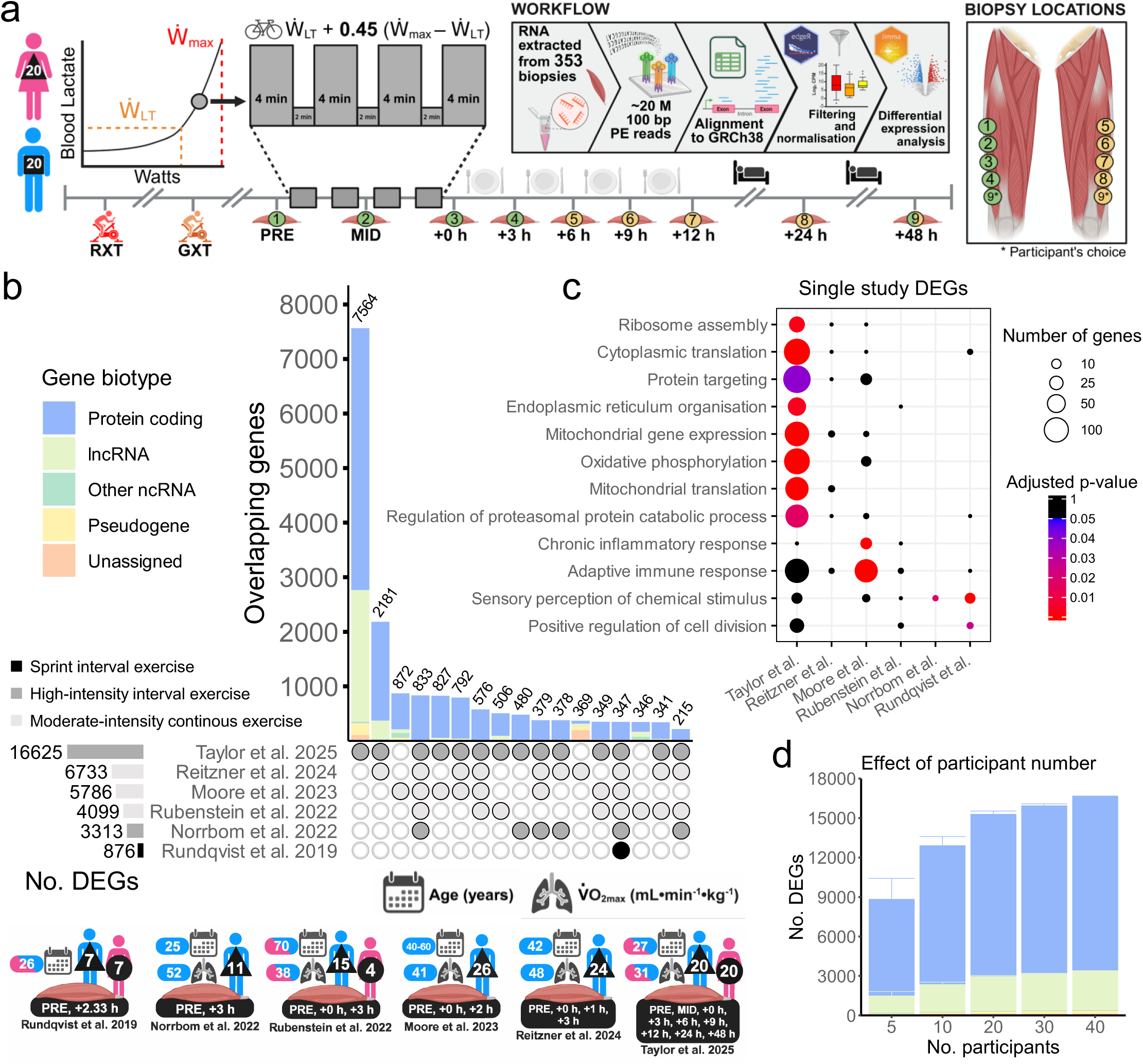
An emphasis on early post-exercise time points has prevented the detection of numerous exercise-induced genes in muscle. **a**Study design, including baseline exercise testing using a ramped exercise (RXT) and graded exercise test (GXT) to determine the peak power output (Ẇmax) and power at lactate threshold (Ẇ_LT_) respectively, prescription of a single session of high-intensity interval exercise (HIIE) using the Ẇmax and Ẇ_LT_, muscle biopsy collection sequence, sleep and meal timing, and bioinformatics workflow. **b** Upset plot comparing the number of differentially expressed genes (DEGs) relative to the PRE time point detected within this study to a selection of previous studies investigating the transcriptomic response to a single session of cardiorespiratory exercise (differential expression was determined by an adjusted p-value < 0.05 using the bioinformatic pipeline described in each study). Participant age, cardiorespiratory fitness, sample size, and the exercise modality used for each study is displayed below. Minimum set size displayed is 200. lncRNA: long non-coding RNA. ncRNA: non-coding RNA. **c** Gene ontology biological process (GOBP) enrichment of DEGs detected within singular studies investigating the transcriptomic response to a single session of cardiorespiratory exercise, significance determined by an adjusted p-value < 0.05 (Benjamini-Hochberg). **d** The number of DEGs (mean ± SD) detected within this study when using a varying number of participants (100 iterations). For each iteration, an equal number of male and female participants were randomly selected. When the total number of participants was 5, the number of males and females was randomly assigned as 2 or 3.

Following transcriptomics data pre-processing (see **Methods**), 17,345 genes remained for further analyses. This included 73% of the skeletal muscle transcriptome annotated within the encyclopaedia of DNA elements (ENCODE) consortium database (**Extended Data Fig. 1a**) and includes 12,312 (95%) of the 12,945 protein-encoding transcripts annotated within the Human Protein Atlas (**Extended Data Fig. 1b**). Consistent with previous studies^13^, the inter-individual variability of gene expression increased in response to exercise, with the median CV increasing from 23% at the PRE time point to a peak of 38% at the +48 h time point (**Extended Data Fig. 1c**), indicating differences in exercise-induced gene expression responses between participants. Following filtering and normalisation, multidimensional scaling revealed separate clusters for males and females, with similar positioning of time-point clusters across both sexes (**Extended Data Fig. 1d**).

In response to exercise, 16,625 genes (96% of all analysed genes) were identified as differentially expressed at one or more time points relative to baseline (PRE) when males and females were analysed together as one cohort. These 16,625 differentially expressed genes (DEGs) included 81% of all previously reported exercised-induced DEGs, as well as more than 7,000 genes not previously reported as responsive to exercise^7–13,23,24^ (**Fig. 1b**; see **Supplementary Data 1 – Tab 1** for a full list of DEGs identified in this study and previous studies). Likely reasons for not detecting all previous exercise-induced DEGs in our study include differences in exercise intensity^7,25^, cardiorespiratory fitness^12,13,26,27^ or health status of the enrolled participants^23^, and RNA isolation and sequencing methodologies used between studies. For the novel exercise-induced DEGs detected within this study, an enrichment was observed for genes involved in numerous gene ontology biological processes (GOBP) – including cytoplasmic translation, mitochondrial gene expression, and protein targeting (**Fig. 1c**; see **Supplementary Data 1 – Tab 2** for a full list of enriched GOBP terms within each study).

The large number of DEGs identified in this study far exceeds the number of genes previously identified as being altered by exercise in two published meta-analyses^23,24^. This likely reflects the unprecedented temporal resolution of our sample collection and the high level of experimental control, as the increased participant number compared to previous studies accounted for less than 15% of the novel exercise-induced genes identified (**Fig. 1d**). Our dataset highlights that much of the exercise transcriptome has been previously overlooked or mischaracterised and will serve as a critical resource, easily accessible via an interactive web application, for understanding and exploring the molecular mechanisms that underlie the many beneficial effects of exercise.

### Exercise-induced gene expression is temporal and persists until at least 48 hours post-exercise

In the present study, the number of upregulated or downregulated genes increased at each consecutive time point until +24 h, followed by the number of DEGs in both directions decreasing slightly at +48 h (**Fig. 2a**; see **Supplementary Data 2 – Tab 1** for the differential expression of each gene in response to exercise). Upregulated genes exhibited a greater average log₂ fold change compared to downregulated genes, as reflected by a more prominent right-skewed log_2_ fold change density distribution (**Fig. 2b**). This asymmetry progressively diminished from the MID to +24 h time point, until ultimately reversing at +48 h. The patterns of differential gene expression were highly variable, with individual genes showing peak log₂ fold changes relative to PRE at various time points, including at +24 h and +48 h post-exercise (**Fig. 2c**).

**Fig. 2:**
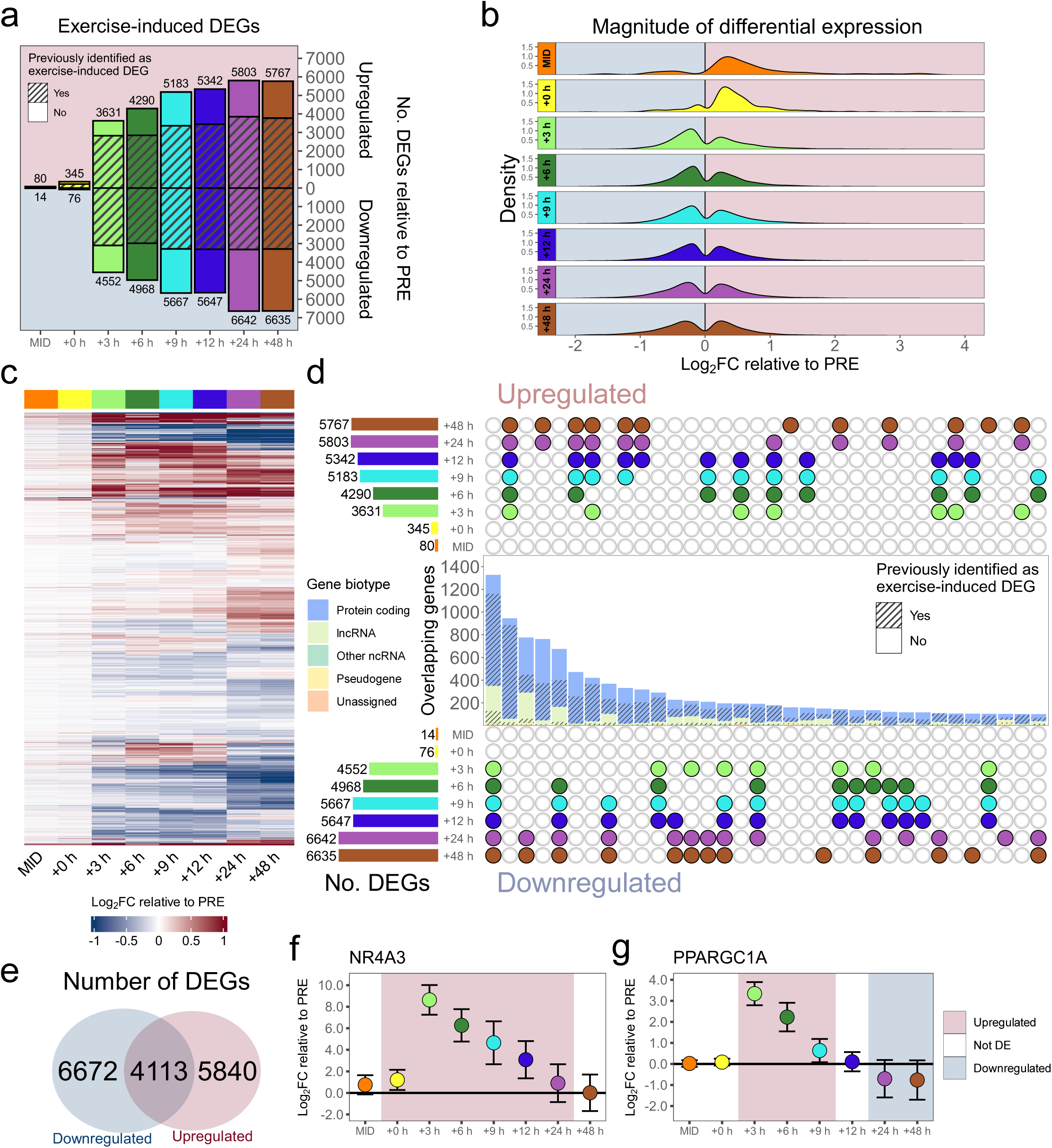
Exercise-induced gene expression is temporal and persists until at least 48 hours post-exercise. **a**The number of upregulated and downregulated differentially expressed genes (DEGs) relative to PRE at each subsequent time point, determined with an adjusted p-value < 0.05 (Benjamini-Hochberg). The number of DEGs with novel and previously identified exercise-induced expression is indicated. **b** Ridge plots illustrating the log_2_ fold change (FC) densities of DEGs at each time point relative to PRE. **c** Heatmap of DEGs at each time point relative to the PRE time point. Row clustering determined by unsupervised hierarchical cluster analysis. **d** Upset plot of upregulated and downregulated DEGs at each time point relative to PRE. The number of DEGs with novel and previously identified exercise-induced expression is indicated for each biotype within an intersection. Minimum set size displayed is 100. lncRNA: long non-coding RNA. ncRNA: non-coding RNA. **e** Venn diagram of the number of DEGs that undergo upregulation or downregulation, or both at different time points, in response to exercise. **f** Differential expression relative to PRE of *NR4A3*. **g** Differential expression relative to PRE of *PPARGC1A*. DE: differentially expressed.

The greatest number of newly identified exercise-induced genes in this study were those upregulated or downregulated at +24 and +48 h post-exercise, whereas fewer previously unreported DEGs were observed at +3 h (**Fig. 2d**). Indicative of a strong focus on upregulated pathways by previous research, a higher proportion of known exercise-induced DEGs were identified in gene sets that were exclusively upregulated, particularly for long non-coding RNA (lncRNA), even though each time point from +3 to +48 h had more downregulated than upregulated genes. This is an intriguing observation, as most studies typically associate exercise with increased expression of specific genes (and their encoded proteins), rather than as a stimulus that also decreases the expression of many genes and impacts nearly all genes expressed in skeletal muscle.

In addition to identifying sets of genes only differentially expressed at specific time points, we observed more than 4,100 genes that were both upregulated and downregulated in response to exercise, depending on the time point investigated (**Fig. 2e**). Consistent with the meta-analysis of Pillon *et al.*^23^, *NR4A3* was the most upregulated gene (as assessed by log_2_ fold change) in response to exercise and followed a ‘canonical’ exercise-induced response^5,6^, with a peak in upregulation at +3 h followed by a return to baseline expression at +48 h (**Fig. 2f**). However, other highly studied exercise-responsive genes, such as *PPARGC1A*, which encodes the protein peroxisome proliferator-activated receptor gamma coactivator 1-alpha (PGC-1α) – a commonly studied regulator of mitochondrial biogenesis, did not follow this pattern (**Fig. 2g**). Consistent with the meta-analysis of Amar *et al.*^24^, *PPARGC1A* was upregulated +3 to +9 h post-exercise but was subsequently not differentially expressed at +12 h and then downregulated at +24 and +48 h post-exercise. This result highlights the value of our approach, as the comprehensive time course of post-exercise muscle biopsies demonstrates how conclusions can vary depending on biopsy timing. This enables, for the first time, consistent tracking of exercise-induced gene expression changes, and allows a deeper investigation into the influence of factors such as cardiorespiratory fitness and biological sex on exercise-induced gene expression.

### Higher cardiorespiratory fitness blunts the exercise-induced expression of genes encoding mitochondrial and ribosomal proteins

In addition to following a time-specific choreography, exercise-induced gene expression has also been shown to be regulated by cardiorespiratory fitness^12,13,26,27^. However, previous studies characterising only early post-exercise responses may have contributed to an underestimation of the impact of cardiorespiratory fitness on exercise-induced gene expression. The range of V̇O_2max_ values in the present study (ranging from 18.4 to 40.0 mL•min^-1^•kg^-1^) (**Fig. 3a**), combined with the large sample size and comprehensive time course of biopsies, has enabled us to conduct the most extensive analysis to date of the association between cardiorespiratory fitness and exercise-induced gene expression.

**Fig. 3:**
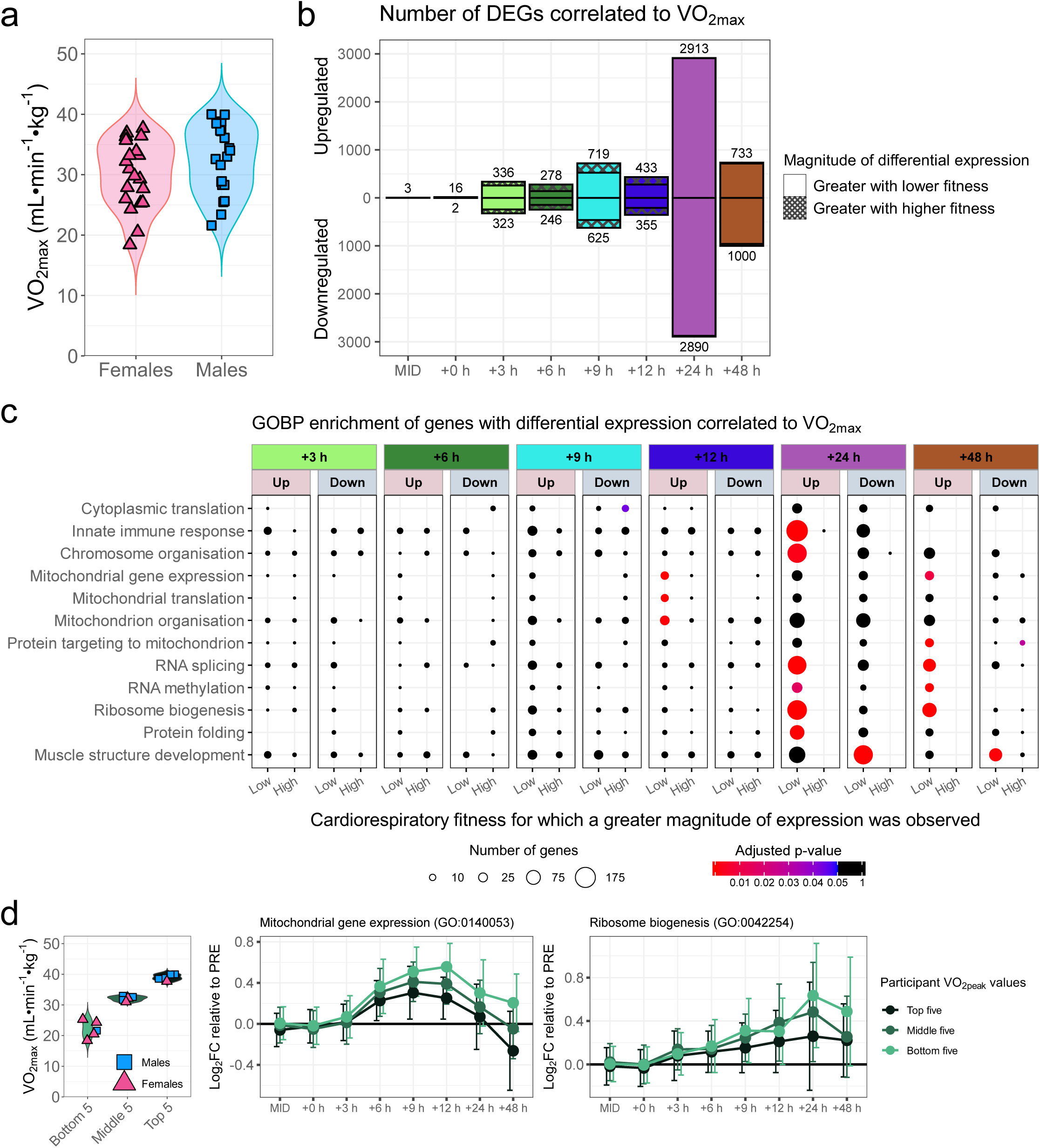
Higher cardiorespiratory fitness blunts the magnitude of exercise-induced expression of genes encoding mitochondrial and ribosomal proteins. **a**Violin plot of V̇O_2max_ values for males and females enrolled within this study**. b** Number of differentially expressed genes (DEGs) at each time point with a log_2_ fold change (FC) relative to PRE correlated with V̇O_2max_ by linear correlation analysis, determined as a p-value < 0.05. **c** Gene ontology biological process (GOBP) enrichment of DEGs correlated to V̇O_2max_ at each time point relative to PRE, significant enrichment determined with an adjusted p-value < 0.05 (Benjamini-Hochberg). **d** Visualisation of the effect of cardiorespiratory fitness on mitochondrial gene expression and ribosome biogenesis using upregulated genes correlated to V̇O_2max_ at +12 h and +24 h respectively. To create separate groups, participants with the five highest, middle, and lowest V̇O_2max_ values were selected and the average expression for all genes in each group was visualised as mean ± SD.

To investigate the effect of cardiorespiratory fitness, each participant’s V̇O_2max_ value was first correlated with the log_2_ fold change relative to PRE of each DEG at each time point (**Fig. 3b**; see **Supplementary Data 3 – Tab 1** for a full list of genes with differential expression correlated to V̇O_2max_). The number of DEGs with differential expression correlated with V̇O_2max_ increased from 3 at MID to 5,803 at +24 h, before decreasing to 1,733 at +48 h. Most of these correlated genes, particularly those at +24 and +48 h, exhibited greater upregulation or downregulation in individuals with lower cardiorespiratory fitness, consistent with a higher cardiorespiratory fitness attenuating exercise-induced changes in gene expression^12,13,26,27^. Given cardiorespiratory fitness may influence patterns of gene expression^28^, such as changing the time at which a gene reaches peak differential expression, but not the peak log_2_ fold change itself, correlations between V̇O_2max_ and the peak upregulation, peak downregulation, standard deviation, and the absolute area under the curve (AUC) for each DEG were also calculated (**Extended Data Fig. 2a**). This revealed an additional 1,488 and 538 genes, with respective correlations observed between V̇O_2max_ and the standard deviation or AUC of gene expression but not the log_2_ fold change at any single time point.

To identify potential exercise-induced gene expression pathways that may be affected by cardiorespiratory fitness, enrichment analysis using the GOBP database was performed for each time point (**Fig. 3c**; see **Supplementary Data 3 – Tab 2** for a full list of GOBP terms correlated to V̇O_2max_ at each time point) and each of the four gene features (**Extended Data Fig. 2b**; see **Supplementary Data 3 – Tab 3** for a full list of GOBP terms correlated to V̇O_2max_ for each gene feature). For the genes with differential expression correlated to V̇O_2max_ at +24 and +48 h, the most significant observed enrichment was for ribosomal biogenesis.

There was a greater differential expression of ribosomal genes when cardiorespiratory fitness was lower (**Fig. 3d**), which supports the hypothesis that exercise induces a shift to a greater capacity for protein translation as cardiorespiratory fitness increases^17^. This may help explain why changes in gene expression in response to a single session of exercise correlate poorly with changes in protein content after training, with much of the regulation of exercise-induced adaptation occurring at the post-transcriptional level^5,6,17^. Further evidence of this is the greater downregulation of genes involved in cytoplasmic translation at +9 h in individuals with higher cardiorespiratory fitness (**Fig. 3c**). As such, once the necessary translation machinery is established, less gene upregulation may be needed following exercise, as observed by the widespread downregulation of gene expression pathways in the skeletal muscle transcriptome of endurance-trained individuals^19^.

Consistent with a lower cardiorespiratory fitness leading to greater improvements in mitochondrial content and respiratory function following training^29^, various terms related to mitochondrial gene expression (**Fig. 3d**), translation, and organisation were significantly upregulated at +12 h in those with a lower cardiorespiratory fitness. Additionally, genes associated with protein targeting to the mitochondria were identified as significantly upregulated in those with a lower cardiorespiratory fitness +48 h post-exercise, indicating specific mitochondrial adaptations may be more prominent at different time points post-exercise. The inclusion of muscle biopsies beyond the previously assessed post-exercise time points has therefore provided a new understanding of the extent to which the exercise-induced expression of genes encoding ribosomal and mitochondrial proteins is blunted in those with a higher cardiorespiratory fitness.

### Mitochondrial gene expression is primarily characterised by downregulation 24 and 48 hours following exercise

As is often depicted in textbooks and reviews, transient increases in the expression of mitochondrial genes have been proposed to underlie subsequent increases in the abundance of mitochondrial proteins following exercise training (**Fig. 4a**)^1,30–33^. Although there is increasing evidence that challenges this dogma^5,6^, our findings highlight that the widespread use of muscle biopsies within only the first few hours post-exercise has meant that hundreds of exercise-induced mitochondrial DEGs have previously evaded detection (**Fig. 1c**), preventing a thorough understanding of exercise-induced mitochondrial gene expression.

**Fig. 4:**
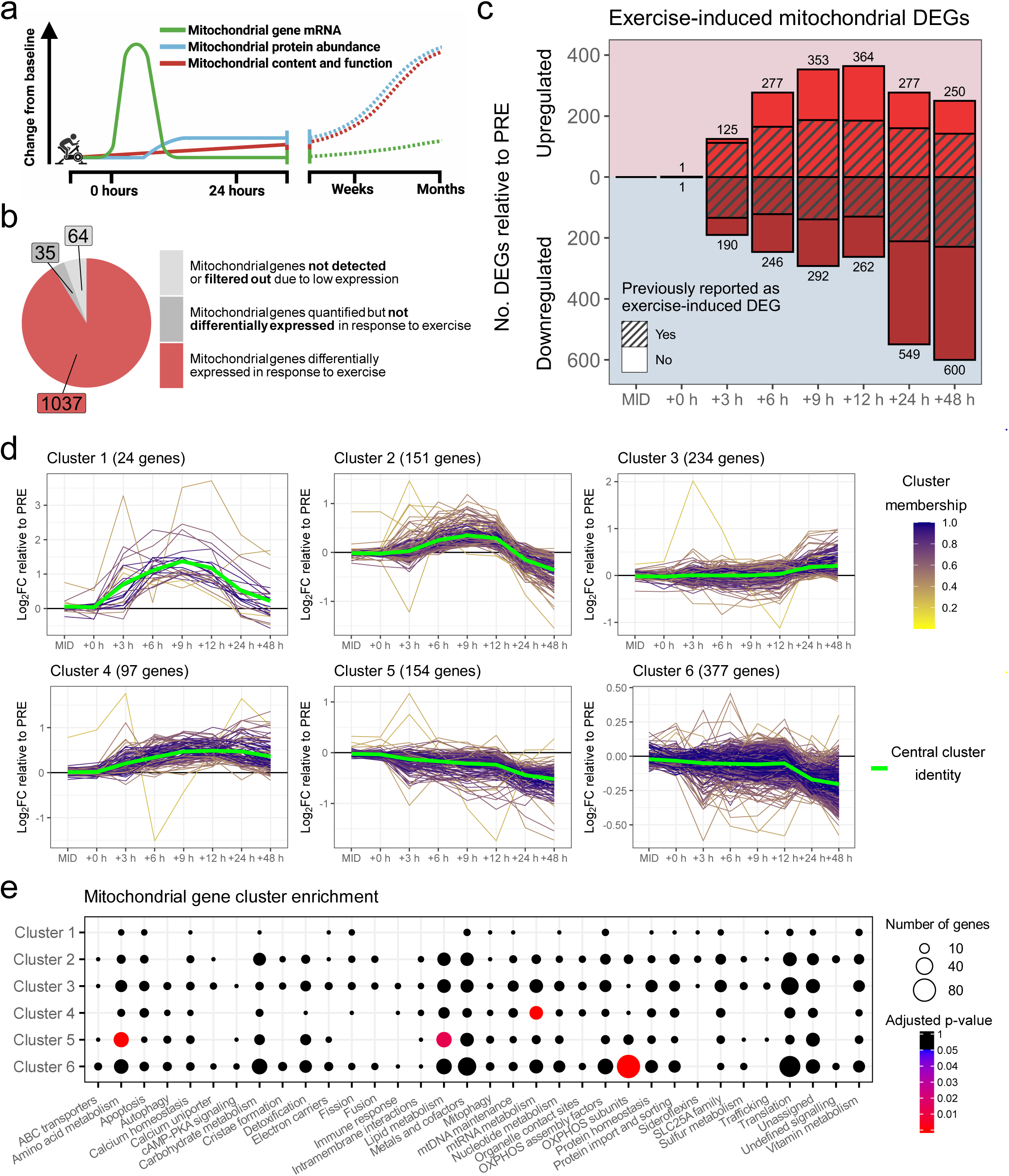
Mitochondrial gene expression is primarily characterised by downregulation 24 and 48 hours following exercise. **a**The prevailing model of exercise-induced mitochondrial biogenesis proposes that a single exercise session triggers a transient increase in transcription factors and coactivators. This leads to the activation of nuclear and mitochondrial gene expression, which promotes the synthesis of mitochondrial proteins. Over repeated exercise sessions, the accumulation of these proteins drives a gradual increase in mitochondrial content and function. **b** Pie plot of the number of mitochondrial differentially expressed genes (DEGs) relative to the total number of proteins with hypothesised or known localisation to the mitochondria (MitoCarta 3.0). **c** Plot of the number of mitochondrial DEGs upregulated and downregulated relative to the PRE time point. The number of DEGs with novel and previously identified exercise-induced expression is indicated. **d** Plots of mitochondrial gene clusters determined using *mfuzz*. The green line represents the cluster’s central expression pattern and other lines the expression of individual mitochondrial DEGs, with colour their membership score – a rating of the similarity of each gene to the central cluster expression pattern. FC: fold change. **e** MitoCarta3.0 pathway enrichment analysis for mitochondrial gene clusters determined using *mfuzz*, significant enrichment determined with an adjusted p-value < 0.05 (Benjamini-Hochberg).

In the present study, 1,037 (91% of the 1,136 genes annotated within MitoCarta3.0) were identified as differentially expressed following exercise (**Fig. 4b**; see **Supplementary Data 4 – Tab 1** for the differential expression of each mitochondrial gene in response to exercise); this includes 565 (50% of all mitochondrial genes) never previously identified as differentially expressed in response to exercise (**Fig. 4c**). The number of upregulated mitochondrial DEGs peaked at +12 h before declining at +24 and +48 h, while the number of downregulated mitochondrial DEGs peaked at +24 and +48 h, encompassing the majority of detected mitochondrial DEGs.

To investigate whether there were specific patterns of expression for mitochondrial DEGs, soft clustering was performed. Using repeated soft clustering (**Extended Data Fig. 3a**) and the visual differentiation of gene expression patterns, six distinctive patterns of mitochondrial gene expression in response to exercise were identified (**Fig. 4d**; see **Supplementary Data 4 – Tab 2** for a full list of mitochondrial clusters and the membership scores for each gene and **Extended Data Fig. 3b** for principal component analysis of cluster centres).

To identify whether specific mitochondrial gene pathways respond similarly following exercise, enrichment analysis of these mitochondrial clusters was performed (**Fig. 4e**). Although significant enrichments were observed for mtRNA metabolism in cluster 4, amino acid metabolism and lipid metabolism in cluster 5, and oxidative phosphorylation (OXPHOS) subunits in cluster 6, all mitochondrial pathways were spread across multiple clusters. The lack of uniformity in differential expression for genes within the same pathway, and even those that require the protein interaction of one another for performing their biological function (e.g., see **Extended Data Fig. 3c** for genes associated with protein import that were found in each of the six clusters), indicate the transcriptional response to exercise is highly varied among mitochondrial genes. Furthermore, the absence of a pronounced increase in the expression of many mitochondrial genes after exercise suggests that post-transcriptional regulation may play an important role in mitochondrial adaptation to exercise. When comparing exercise-induced mitochondrial gene expression to previously reported increases in mitochondrial protein abundance following similar training^27^, no consistent pattern emerged – regardless of cluster identity or direction of gene expression change (**Extended Data Fig. 3d**). These results further challenge the longstanding dogma that proposes a direct relationship between exercise-induced increases in mRNA levels and subsequent changes in the abundance of the proteins they encode^5,6^.

### Minimal effect of sex on exercise-induced gene expression

Although sex-specific differences in skeletal muscle structure and exercise metabolism have been reported^16^, many studies investigating differential gene expression between males and females have used cohorts not matched for age or cardiorespiratory fitness. This is problematic as previous studies have observed changes in gene expression with ageing^17^, and we and others^12,13,26,27^ have reported an effect of cardiorespiratory fitness on exercise-induced gene expression (**Fig. 3b**). While the well-documented differences between males and have been used to support the notion of a sex-specific transcriptional response to exercise, empirical support for this hypothesis is lacking.

Consistent with previous research indicating sex-specific differences in baseline skeletal muscle gene expression using RNA-seq^17–22^, 1,193 genes (6.9% of all quantified genes) were differentially expressed between males and females at the PRE time point (see **Supplementary Data 5 – Tab 1** for the full list of genes with sex-specific differential expression at baseline). An overrepresentation of DEGs were from the X (10.6% of all X-encoded genes detected) and Y chromosomes (100% of all Y-encoded genes detected), consistent with their roles in sexual differentiation (**Extended Data Fig. 4a**). However, no other clear sex-specific patterns of expression were detected, with enrichment analysis only identifying significantly enriched terms related to urogenital development for the 675 genes upregulated in males or the 518 genes upregulated in females (**Extended Data Fig. 4b**; see **Supplementary Data 5 – Tab 2** for a full list of GOBP terms enriched for sex-specific differential expression at baseline). Although most differences in sex-specific expression at baseline between this study and previous studies using RNA-seq are congruent, with only a small number of genes upregulated in opposing sexes between studies^17–22^ (**Extended Data Fig. 4c**; see **Supplementary Data 5 – Tab 3** for a full list of sex-specific DEGs at baseline identified in this study and previous studies), very little overlap of genes with sex-specific expression at baseline, excluding those encoded on the Y chromosome, is observed between studies (**Extended Data Fig. 4d**). The limited enrichment of sex-specific pathways in our study, compared to previous ones, may reflect our close matching for age and cardiorespiratory fitness, in contrast to earlier studies that used tissue biobanks or cohorts where participant exercise habits were not recorded.

In response to exercise, an overlap of 14,662 exercise-induced DEGs relative to PRE was observed between sexes, with a further 810 genes differentially expressed only within males (5.2% of the 15,472 DEGs in males) and a further 987 genes differentially expressed only within females (6.3% of the 15,649 DEGs in females) (see **Supplementary Data 5 – Tab 4** and **Supplementary Data 5 – Tab 5** for the sex-specific differential expression in response to exercise for females and males respectively). Although the number of sex-specific and shared DEGs at each time point fluctuated between males and females, the proportion of previously identified to novel exercise-induced genes within both sexes was comparable at each time point (**Fig. 5a**). Visualisation of the average log_2_ fold change for shared and sex-specific DEGs indicated that genes identified as differentially expressed in only males or females had on average a lower differential expression than those DEGs that were common to both sexes at each time point (**Fig. 5b**). This suggests that differences in DEGs identified between males and females may have arisen due to the lower statistical power from using 20 participants of each sex per group^34^.

**Fig. 5:**
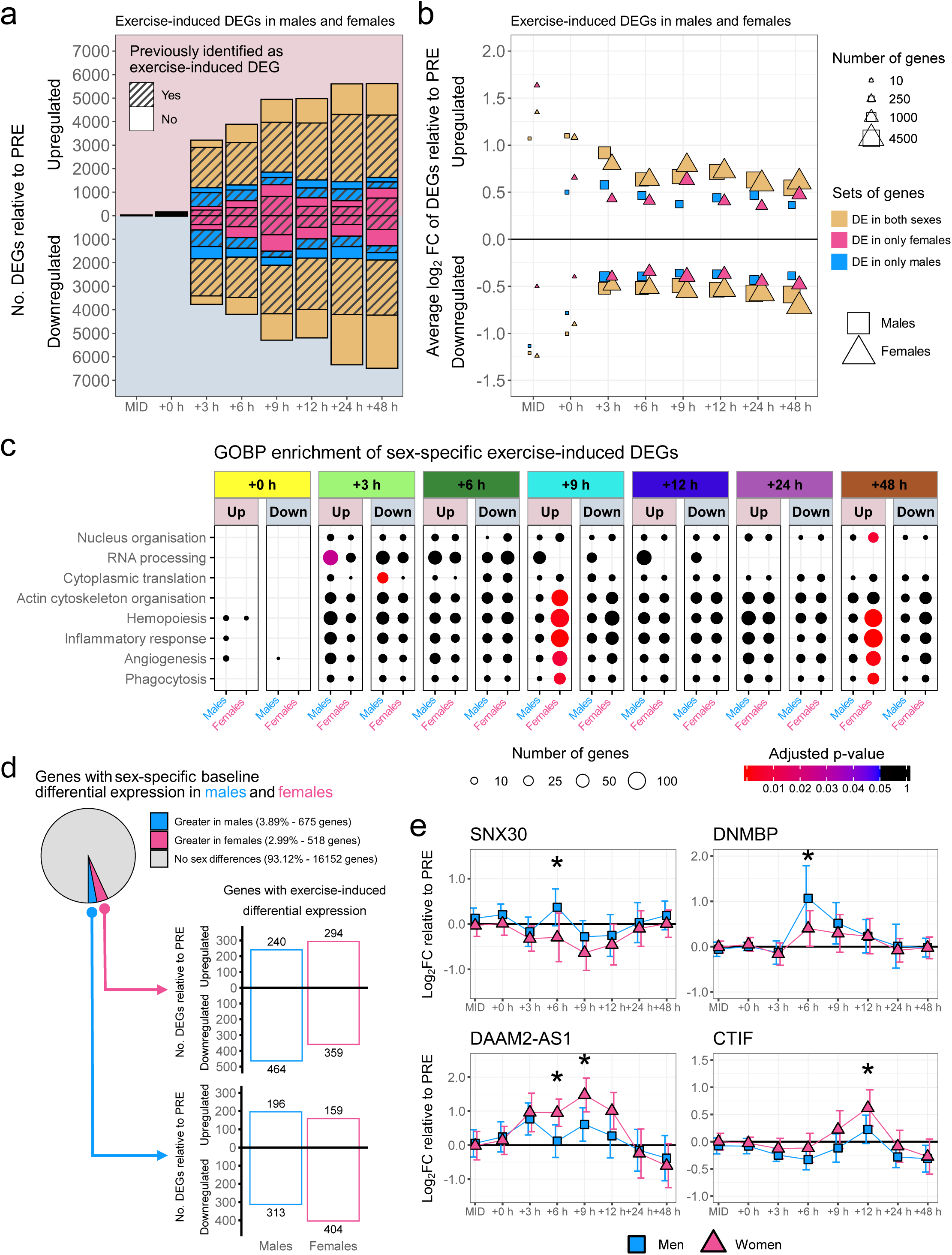
Minimal effect of sex on exercise-induced gene expression. **a**The number of upregulated and downregulated differentially expressed genes (DEGs) relative to PRE at each subsequent time point, determined with an adjusted p-value < 0.05 (Benjamini-Hochberg (BH)), within males and females. The number of DEGs with novel and previously identified exercise-induced expression is indicated. **b** Plot of the average log_2_ fold change (FC) of DEGs within males and females for each time point relative to PRE. DE: differentially expressed. **c** Gene ontology biological process (GOBP) enrichment for genes with significant differential expression in response to exercise within only males or females, significant enrichment determined with an adjusted p-value < 0.05 (BH). **d** The number of upregulated and downregulated DEGs across all time points relative to PRE in males and females for genes with sex-specific differential expression at baseline. **e** The four DEGs, *SNX30*, *DNMBP*, *DAAM2-AS1*, and *CTIF*, with sex-specific differential expression relative to PRE, determined with an adjusted p-value < 0.05 (BH). Significance for a specific time point relative to PRE indicated by *.

To identify whether specific pathways were upregulated or downregulated to a greater extent in males or females, enrichment analysis using the GOBP database was performed (**Fig. 5c**; see **Supplementary Data 5 – Tab 6** for a full list of GOBP terms enriched for sex-specific differential expression in response to exercise). Although several pathways showed a greater number of DEGs at specific time points in males and females, many of these enrichments involved upregulated processes related to inflammation and circulation and were confined to the +9 and +48 h biopsies. These differences may reflect slight differences in cell type composition within the muscle biopsies collected from each sex. Supporting this interpretation, the proportion of sex-specific DEGs for these enriched pathways was remarkably similar at other time points.

Although our previous observations indicate minimal sex-specific differences, genes with higher baseline expression in one sex trended towards being downregulated in that same sex and upregulated in the opposite sex following exercise (**Fig. 5d**). That is, genes with higher baseline expression in females trended towards being more frequently upregulated in males and downregulated in females following exercise, and vice versa for genes with higher baseline expression in males. While speculative, these findings suggest a potential attenuation of sex-specific differences in skeletal muscle following exercise training. Previous studies have shown that baseline expression differences between males and females diminish as cardiorespiratory fitness increases^19^, a trend that has also been reported for protein abundance^35^.

Nevertheless, when directly comparing exercise-induced differential expression relative to PRE between sexes, only four (*SNX30*, *DNMBP*, *DAAM2-AS1*, and *CTIF*) of the 17,345 quantified genes were significantly different at any time point relative to PRE (**Fig. 5e**; see **Supplementary Data 5 – Tab 7** for differences in differential expression between each sex at each time point). *SNX30*, part of the sorting nexin family involved in protein trafficking, was previously reported to be more highly expressed in females at baseline^22^, a finding also observed in this study. *DNMBP*, which encodes a dynamin-binding guanine nucleotide exchange factor, has shown higher baseline expression in females in one previous study^20^. *DAAM2-AS1*, the antisense strand of a gene involved in Wnt signalling gene, was more highly expressed in males in this study as well as one previous study^20^, however, its function remains uncharacterised. *CTIF*, a component of the CBP80 translation initiation complex, has never been identified as having sex-specific expression at baseline. None of these genes have previously shown sex-specific differential expression from exercise^21,36^, or sex-specific protein abundance at baseline^21,35,37^ or following training^21,37^. Compared to the differential expression of individual time points, analysis of the area under the curve for all differentially expressed genes revealed no significant differences between sexes, further challenging the accepted dogma of meaningful sex-specific differences in exercise-induced gene expression.

The limited sex-specific differences observed in the present study are likely attributable to the careful matching of males and females for cardiorespiratory fitness and relative exercise intensity. This is underscored by the finding that approximately 60% of all DEGs identified in this study exhibited exercise-induced differential expression that was correlated with V̇O_2max_ at one or more time points or to a gene feature. Additionally, by setting exercise intensity relative to Ẇ_LT_ and Ẇ_max_, rather than as a fixed percentage of Ẇ_max_, HR_max_, or V̇O_2max_, a more comparable metabolic stress across individuals with varying fitness levels^38^ and between sexes^39^ was achieved. To determine if the lack of sex-specific differences in exercise-induced gene expression translates to similar training adaptations, additional studies appropriately matching males and females for cardiorespiratory fitness and exercise stimulus are required.

## Discussion

Our greater coverage of the post-exercise transcriptional response allowed us to identify more than 16,600 differentially expressed genes. Of these genes, more than 7,000 have not previously been identified as differentially expressed in response to exercise. The magnitude of differential expression for numerous pathways was influenced by cardiorespiratory fitness, with these pathways enriched for genes encoding mitochondrial and ribosomal proteins at various time points post-exercise. Many of the differentially expressed mitochondrial genes were characterised only by downregulation in response to exercise, indicative of a greater influence of post-transcriptional regulation than previously appreciated. Despite there being 1,193 differentially expressed genes between males and females at baseline, minimal sex-specific differences were observed for exercise-induced gene expression.

The lack of a non-exercise control group prevented the identification of genes with uncharacterised circadian patterns, nor assess the effect of exercise on circadian gene expression. However, consistent with prior studies, upstream circadian regulators *CLOCK* and *BMAL1* returned to baseline by +24 and +48 h post-exercise (**Extended Data Fig. 5a**)^40^. In contrast, downstream regulators *PER1* and *CRY1* showed differential expression at these time points, suggesting a phase shift aligned with exercise timing, with peak expression shifted toward the morning compared to non-exercising controls from previous circadian assessments (**Extended Data Fig. 5b**)^41^. In studies investigating exercise-induced gene expression that have included non-exercise controls, only a small fraction of skeletal muscle genes has shown circadian expression^42^. However, whether DEGs in non-exercise controls truly have circadian rhythm cannot be discerned, as, like this study, the potential local and systemic inflammatory effects of repeated muscle biopsies on gene expression cannot be completely excluded^43^. While a strength of our study is the large number of biopsies collected from each participant, it is possible that damage and inflammation from each biopsy may provoke a transcriptional response. This remains a controversial topic, with some studies reporting changes in mRNA with repeated muscle biopsy sampling^43^, especially for inflammation-related genes^44^.

However, genes involved in other adaptive processes, such as mitochondrial biogenesis, show little or no change when biopsies are taken from separate incisions and sections of the same muscle belly^45,46^, as performed in this study. Switching biopsy sites from the first to the second leg (from the +3 h to the +6 h biopsy) was associated with a small decrease in the magnitude of differential expression in the GOBP term ‘regulation of skeletal muscle regeneration’ (**Extended Data Fig. 6a**). However, few of the genes in this GOBP term that overlapped between +3 h and +9 h were not also differentially expressed at +6 h (**Extended Data Fig. 6b**), with exceptions being the muscle repair gene *KLF5* and extracellular matrix component *COL6A1* (**Extended Data Fig. 6c**). However, pathways unrelated to inflammation appear unaffected by switching the biopsy site from the first to the second leg, as shown by the stable changes in the number of DEGs between time points (**Fig. 2a**), the distribution of log_2_ fold changes (**Fig. 2b**), and in pathways like mitochondrial gene expression (**Fig. 4c**).

Since RNA-seq was performed on homogenised whole-muscle, no direct quantification of the specific contribution from different cell types present in each biopsy can be discerned^47^. Nonetheless, when performing cell type deconvolution using the Tabula Sapiens single-cell RNA-seq (scRNA-seq) atlas to estimate the presence of various cell types in each biopsy, we observed differences in the number of detected cell types across time points (**Extended Data Fig. 7a**). However, when examining the contribution of each cell type, most of the RNA was identified as originating from fast and slow muscle cells (**Extended Data Fig. 7b**).

Although several less abundant cell types showed changes relative to PRE at various time points in both males and females, no significant sex differences were observed in these relative abundance changes. Despite this, sex-specific differences within time points were observed for fast muscle cells at +9 h and slow muscle cells at +24 h and +48 h, with females showing lower proportions compared to males for each. As such, these differences may have contributed to the enrichment of pathways related to inflammation and circulation, which were significantly upregulated exclusively in females, and only at +9 h and +48 h (**Fig. 5c**).

This study recruited healthy, young, untrained males and females and, therefore, some of the findings may not be translatable to individuals of different ages or health status. Although available evidence indicates that the menstrual phase has minimal to no effect on exercise physiology^48^, a lack of standardisation in our participants’ menstrual phase may have also influenced our results. We employed a specific HIIE protocol known to elicit robust increases in various mitochondrial characteristics^27^. However, this could limit the generalisability of our findings to other exercise modalities, such as resistance training, moderate-intensity continuous exercise, or sprint interval training, where distinct adaptations may occur^49^.

To facilitate accessibility of this dataset, we have developed an interactive Shiny web application that allows users to explore exercise-induced gene expression for individual genes or pathways and conduct clustering analyses. Collectively, our findings regarding the temporal response of exercise-induced gene expression will help inform decisions regarding the timing of biopsies for studies investigating the transcriptional response to exercise and help improve the design of future experiments in this continually growing field.

## Methods

### Participants and ethics approval

Twenty healthy males and twenty healthy females volunteered to participate in this study (physiological and performance parameters are presented in **Table 1**). Potential subjects were deemed eligible if aged 18–40 y, were untrained (i.e., < 1.5 h per week of structured aerobic activity for six months prior to the study) and were non-smokers prior to and throughout the study. Participants underwent medical screening to exclude conditions that may have precluded their participation (e.g., cardiovascular, musculoskeletal, pregnancy, and/or metabolic disorders), and were informed of all study requirements, risks, and benefits, before giving written informed consent. Approval for the study’s procedures, which conformed to the standards set by the latest revision of the Declaration of Helsinki, was granted by the Victoria University Human Research Ethics Committee (HRE20-212) and the Australian Catholic University Human Research Ethics Committee (2022-2518R).

**Table 1.** Participant characteristics.

| Measurement | Males | Females |
| --- | --- | --- |
| Age (years) | 27.3 ± 6.3 | 27.6 ± 5.6 |
| Height (cm) | 179.6 ± 10.2 | 166.5 ± 7.8* |
| **Weight (kg)** | 81.5 ± 12.5 | 68.0 ± 9.9* |
| **BMI (kg·m<sup>-2</sup>)** | 25.3 ± 3.8 | 24.5 ± 3.0 |
| **$\dot{V}O_{2\max}$ (L·min<sup>-1</sup>)** | 2.6 ± 0.6 | 2.0 ± 0.3* |
| **$\dot{V}O_{2\max}$ (mL·min<sup>-1</sup>·kg<sup>-1</sup>)** | 32.2 ± 5.7 | 30.1 ± 5.6 |
| **$\dot{W}_{\max}$ (W)** | 244.5 ± 56.6 | 162.4 ± 33.2* |
| **$\dot{W}_{LT}$ (W)** | 130.5 ± 35.4 | 88.5 ± 21.2* |
| **BLa at $\dot{W}_{LT}$ (mmol·L<sup>-1</sup>)** | 3.4 ± 1.0 | 2.9 ± 0.8 |
| **HIIE power (W)** | 179.3 ± 43.5 | 123.2 ± 25.3* |
| **HIIE power (% $\dot{W}_{\max}$)** | 72.8 ± 4.7 | 75.9 ± 4.2* |

BMI: body mass index; $\dot{V}O_{2\max}$ : maximum oxygen uptake; $\dot{W}_{\max}$ : maximum power output; $\dot{W}_{LT}$ : power at lactate threshold; BLa: blood lactate concentration; HIIE: high-intensity interval exercise. \* $p < 0.05$ , significant difference between sexes. All values are mean ± SD.

### Testing procedures

Participants were instructed to refrain from strenuous exercise in the 48 h prior to testing, from alcohol for 24 h prior to testing, and from food and caffeine 3 h prior to testing. All exercise tests were performed at approximately the same time of day for each participant throughout the study to avoid differences caused by potential variations in circadian rhythm or daily fatigue.

#### RXT

Participants completed a ramped exercise test (RXT) to determine their maximum power output (Ẇ_max_) and maximum oxygen uptake (V̇O_2max_). Demographic data, physical activity ratings, height, and body mass data were used to devise a customised test protocol of 1-min stages using an electronically braked cycle ergometer (Lode Excalibur Sport, Groningen, Netherlands), with a desired test duration of 8–12 min^50^. Participants were instructed to cycle between 70 and 85 revolutions per minute (rpm), with the resistance starting at 20% of their estimated Ẇ_max_ before increasing every 1 min until exhaustion (defined as the inability to maintain a cadence above 60).

Expired gases were sampled every 15 s throughout the test using a MOXUS Modular Metabolic System (AEI Technologies, Pennsylvania, USA), and the average of the two highest 15-s readings was recorded as a participant’s V̇O_2max_. After 5 min of rest, participants performed a verification bout to confirm their V̇O_2max_ value. In the verification bout, the power output was set at 105% of Ẇ_max_ reached in the ramp protocol, and the participants cycled to exhaustion without a limit on maximum cadence^50^. Heart rate (Polar Electro Oy, Kempele, Finland) and a Borg rating of perceived exertion (RPE) were recorded in the last 10 s of each stage.

#### GXT

Participants completed a graded exercise test (GXT) to determine the power attained at their lactate threshold (Ẇ_LT_). Before the test, a cannula was inserted into an antecubital vein for subsequent venous blood sampling by a trained phlebotomist or single-use lancing needles (Accu-Chek Safe T-Pro Plus Lancet, Roche, Basel, Switzerland) and capillary tubes were used. The GXT consisted of 4-min exercise stages using an electronically braked cycle ergometer (Lode Excalibur Sport) that progressively increased in resistance by 7.6% until exhaustion (defined as the inability to maintain a cadence above 60), with a desired test duration of 32–40 min. The participants were instructed to cycle at a cadence between 70 rpm and 85 rpm, with the intensity starting at 20% of their estimated peak power (which was estimated using demographic data, physical activity ratings, height, and body mass data, as previously used in our lab^50^).

Blood was sampled at rest and immediately after completion of each 4-min stage, and lactate concentration was determined using an automated analyser (YSI 2500 or YSI 2000 Glucose/Lactate Analyser, YSI, Ohio, USA). The lactate threshold (Ẇ_LT_) was determined by the modified Dmax method (Dmax_MOD_)^50^. Expired gases were sampled every 15 s using a MOXUS Metabolic Cart System (AEI Technologies) and the average of the two highest 15-s readings was recorded as a participant’s V̇O_2max_ for the test. Heart rate (Polar Electro Oy) and a Borg RPE were recorded during the last 10 s of each stage.

### Muscle biopsy and exercise protocol

Participants were instructed to fast 12 h prior to the HIIE session, not to consume any caffeine in the 12 h prior, not to consume any alcohol in the 24 h prior, and to avoid exercise in the 24 h prior. A medical doctor obtained skeletal muscle biopsies (∼100 mg samples) under local anaesthesia (5 mg•mL^-1^ Lidocaine) from the *vastus lateralis* using the suction-modified Bergström muscle biopsy technique.

Following a resting biopsy (PRE), participants began the HIIE session consisting of four 4-min intervals, separated by 2 min of rest, on an electronically braked cycle ergometer (Lode Excalibur Sport). The intensity of each interval was set at Ẇ_LT_ + 0.45 (Ẇ_max_ – Ẇ_LT_). Additional muscle biopsies were collected mid-exercise (MID; between the 2nd and 3rd interval), immediately post-exercise (+0 h), and at +3, +6, +9, +12, +24, and +48 h following completion of the HIIE session. To minimise the influence of local inflammatory responses, each biopsy was taken from a separate incision at least 1 cm distal to the previous site. To control and assess the effect of repeated muscle biopsies, the initial four muscle biopsies were collected from one leg and the subsequent four from the opposite leg, with the leg for the final muscle biopsy chosen based on the participant’s preference. Immediately after being rapidly cleaned of excess blood, fat, and connective tissue, the biopsies were flash frozen using liquid nitrogen for subsequent RNA extraction.

Standardised meals based on the *Australian Dietary Guidelines (2013)* were provided for the 24 h following the HIIE session to minimise any confounding effect of diet. The macronutrient composition of these meals was 50% carbohydrates, 30% fat, and 20% protein. Each participant’s energy intake was calculated using the Mifflin-St Jeor equation, with the activity multiplier of 1.2, and the addition of estimated energy expenditure from the HIIE session.

### RNA-seq analysis

#### RNA extraction

RNA sequencing (RNA-seq) was performed on skeletal muscle biopsies collected before (PRE), during (MID), immediately after (+0 h), as well as +3, +6, +9, +12, +24 and +48 h post exercise. Two participants withdrew due to personal reasons, including one male following the +3 h biopsy and one female following the +12 h biopsy, leaving 353 samples for RNA-seq analysis (**Extended Data Fig. 8**). To perform RNA-seq, RNA was extracted from 10–20 mg of frozen skeletal muscle homogenised using the TissueLyser II (Qiagen, Maryland, USA) and isolated using the RNeasy® Plus Universal Mini Kit (Qiagen) according to manufacturer’s instructions (except for the use of 2-proponal instead of ethanol for isolation of the aqueous phase).

#### RNA sequencing and assembly

Samples were sequenced (100 bp, paired-end reads) using the DNBseq™ platform at the Beijing Genomics Institute (BGI) in Hong Kong, China. Per base sequence quality for all samples was > 90.1% bases above Q30. Transcriptome assembly was completed with reads screened for the presence of any adaptor/over-represented sequences and cross-species contamination using SOAPnuke (v2.1.7), with approximately 20 M clean reads per sample remaining for further analysis. Cleaned sequence reads were aligned against the *Homo sapiens* genome (GRCh38.p14) using STAR aligner (v2.7.11b). Counts of reads mapping to each known gene were summarised using featureCounts (v2.1.1) to provide the matrix used for subsequent analysis.

#### Bioinformatic analysis of RNA-seq data

Count data was filtered to remove lowly expressed genes with average counts per million (CPM) below ten at any time point and then normalised using the method of trimmed mean of M-values (TMM) with the *edgeR* (v.4.4.2) package in R (v.4.4.3). Differential expression analysis was completed with the *edgeR* and *limma* (v.3.62.1) packages in R, with the voomLmFit function used to estimate mean-variance relationships and apply a linear model, while allowing for the loss of residual degrees of freedom for groups with exactly zero counts. The eBayes function was used to subsequently provide empirical Bayes moderation for the model. Upset plots were visualised using the *ComplexUpset* (v.1.3.3) and *ggplot2* (v.3.5.2) packages in R. Gene set enrichment analysis was completed using the *enrichR* (v.3.4) and *clusterProfiler* (v.4.14.6) packages in R. Unsupervised hierarchical clustering with the ‘average’ method for Euclidean distances was used to visualise differentially expressed genes in heatmaps by the package *ComplexHeatmap* (v.2.22.0) in R. Area under the curve (AUC) was calculated by the trapezoidal rule using the *pracma* (v.2.4.4) package in R. Soft clustering of mitochondrial genes was completed using the *mfuzz* (v.2.66.0) package in R. Cell type deconvolution was completed using the *MuSiC* (v.1.0.0) package in R and the Tabula Sapiens database. A male +24 h sample was removed from the analysis due to the lack of skeletal muscle gene expression using cell type deconvolution. Additional plots were visualised using the *ggplot2* and related packages in R.

### Statistical analysis

For transcriptomic analysis, significance was determined by an adjusted p-value < 0.05 using the Benjamini–Hochberg method. Linear correlations were performed using the *rstatix* (v.0.7.2) package in R, with a p-value < 0.05 indicating significance. For cell type deconvolution, linear mixed-effects models were fitted for each cell type using the *lme4* package (v1.1-37) in R, with fixed effects for the interaction between time and sex, and random intercepts for participant ID. Significant effects were further examined using multiple comparisons with the *emmeans* (v1.11.2) package in R, applying a Šidák correction to control the family-wise error rate, with an adjusted p-value < 0.05 indicating significance. For the comparison of physiological data between sexes, Welch’s t-tests were performed using the *stats* (v.4.4.2) package in R, with a p-value < 0.05 indicating significance. All values are reported or visualised as means ± SD, unless otherwise specified. Additional statistical details for each analysis can be found in the associated figure legends.

### EXERgene web application

To improve the accessibility of this dataset, a user-friendly web application was generated using R Shiny (https://shiny.rstudio.com). This application allows selection of single genes or gene ontology (GO) terms to display an interactive graph of differential expression. The application additionally allows the toggling between sexes, levels of cardiorespiratory fitness, and soft clustering of selected genes using *mfuzz*. Code is available at https://github.com/DaleFTaylor.

## Acknowledgements

We thank all the participants for their time and effort in the study. The authors would also like to thank the technical staff at Victoria University for their assistance throughout the study. We also thank the creators of all the repositories, software, and R packages used in this study, whose contributions were invaluable but could not be individually cited due to reference limitations. Figure 1a, 1b, 4a, Extended Data Figure 1d, Extended Data Figure 3c, Extended Data Figure 4d, Extended Data Figure 6a, Extended Data Figure 8 were partially or fully created with BioRender (www.biorender.com) and the relevant licensing rights was obtained. This study was funded by an Australian Research Council (ARC) Discovery Project grant (DP200103542).

## Contributions

D.J.B., J.H., R.B., and N.J.H. conceptualised the study. D.J.B., J.H., N.J.H., and D.F.T. devised the study methodology. D.F.T., E.G.R., and A.G. delivered the exercise sessions and performed sample collection. D.F.T. performed the sample processing. RNA-seq analysis was performed at the Hong Kong Facility of BGI genomics. D.F.T. and N.J.C. performed statistical and bioinformatic analysis. D.F.T. delivered the visualisation. D.F.T. and D.J.B. wrote the initial draft of the manuscript. N.J.H., J.A.H., N.J.C., and D.J.B. provided supervision. D.J.B., J.A.H., R.B., and N.J.H. funded the research. D.F.T. and D.J.B. have primary responsibility for final content. Data collection took place at Victoria University. Muscle analysis took place at Victoria University and BGI genomics. All persons designated as authors qualify for authorship, and all those qualifying for authorship are listed. All authors have read and approved the final manuscript.

---

**Extended Data Fig. 1:**
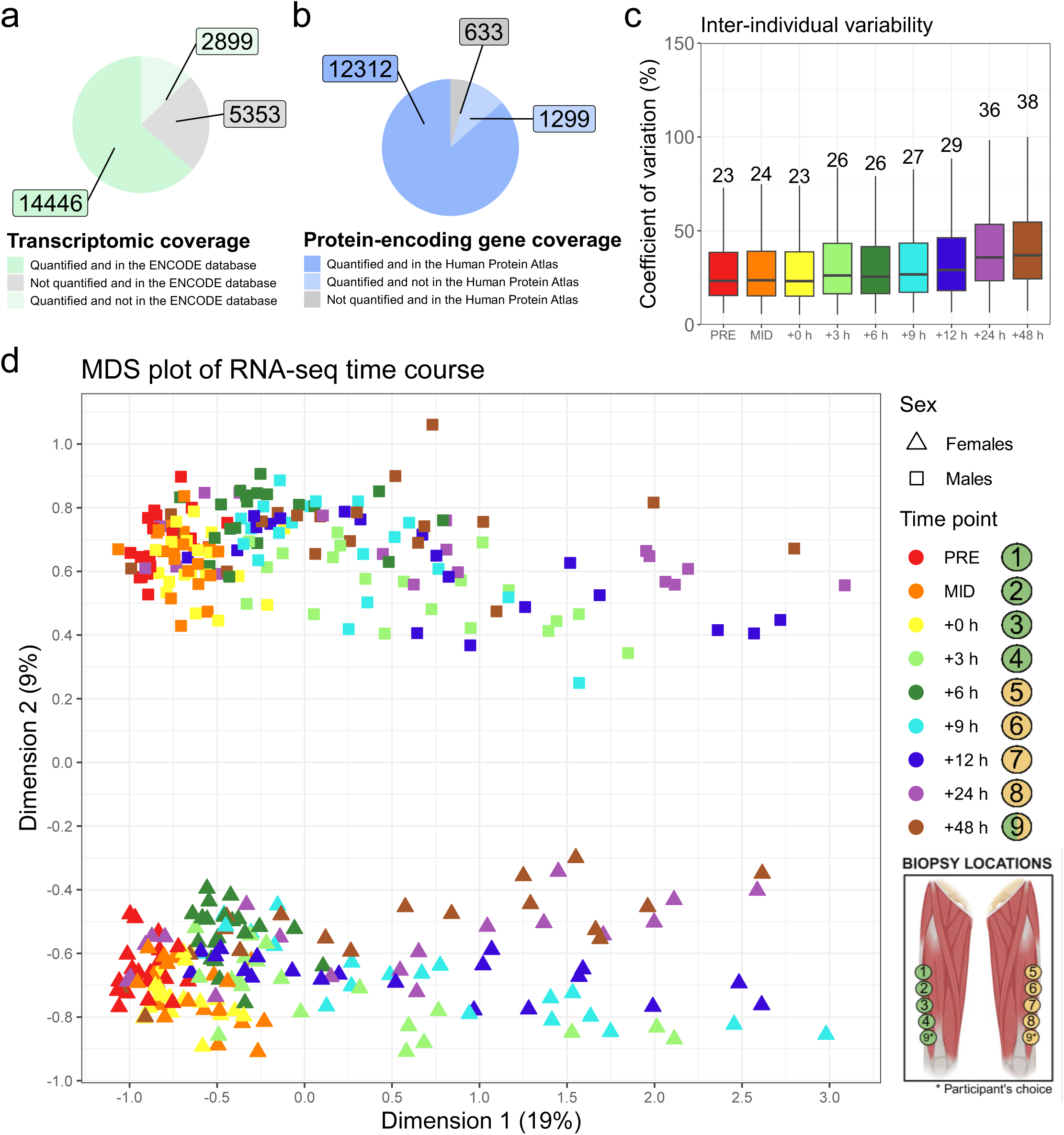
Sample gene coverage, inter-individual variability, and divergence. **a**RNA sequencing (RNA-seq) transcript depth compared to the skeletal muscle database of the ENCODE project. **b** Protein-coding transcript depth compared to the skeletal muscle database of the Human Protein Atlas. **c** Inter-individual variability of all detected genes using raw count per million (CPM) values at each time point. Each box represents the interquartile range (IQR) and median of the coefficient of variation (also displayed as text). Each whisker representing 1.5 * IQR for both directions. Outliers are not displayed. **d** Multidimensional scaling (MDS) of CPM values following filtering and normalisation.

**Extended Data Fig. 2:**
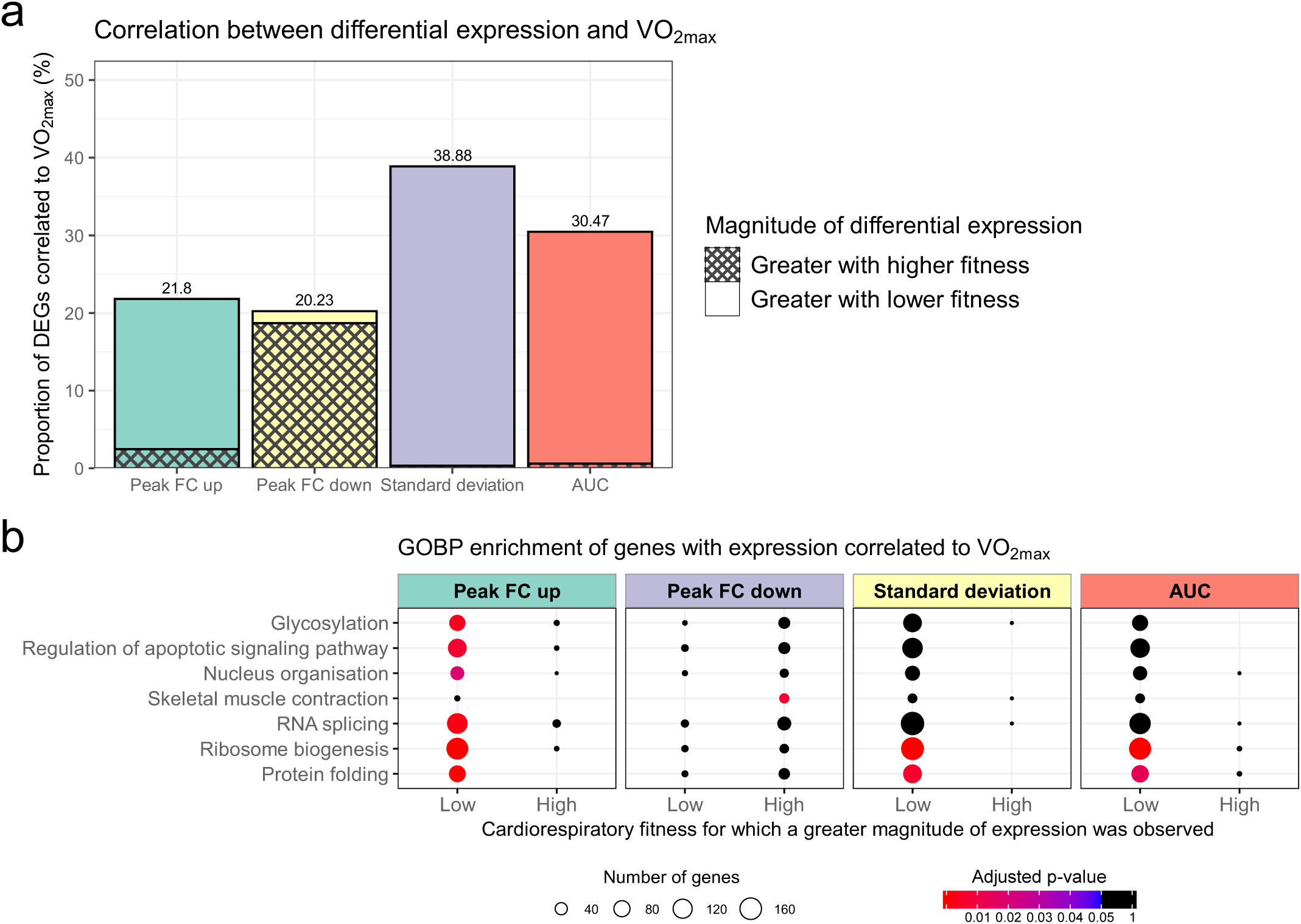
The effect of cardiorespiratory fitness on exercise-induced gene expression (gene features). **a**Proportion of differentially expressed genes (DEGs) with a log_2_ fold change relative to PRE for gene features correlated with V̇O_2max_ by linear correlation analysis, determined as a p-value < 0.05. **b** Gene ontology biological process (GOBP) enrichment of DEGs correlated to V̇O_2max_for gene features, significant enrichment determined with an adjusted p-value < 0.05 (Benjamini-Hochberg).

**Extended Data Fig. 3:**
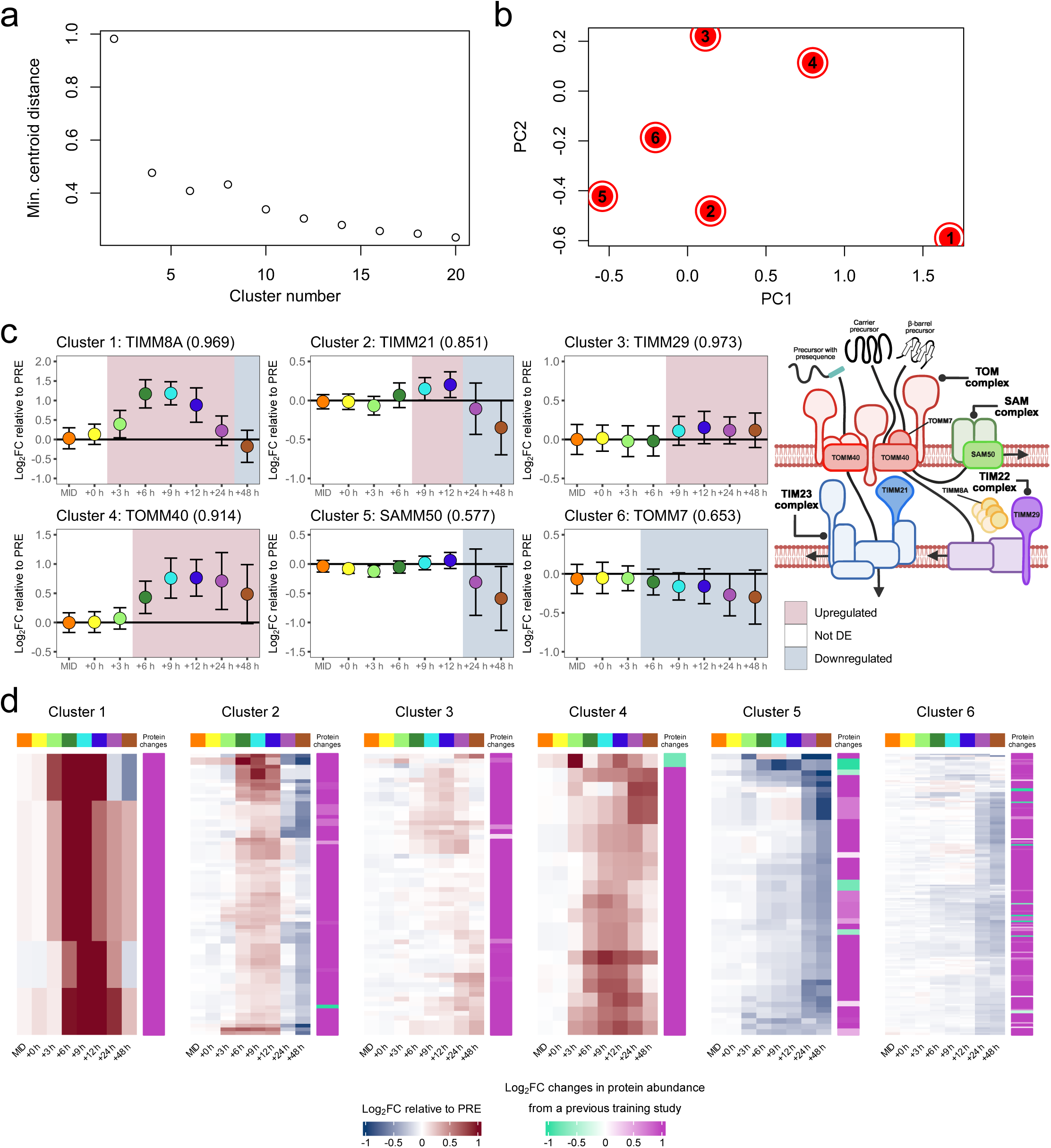
Clustering of differentially expressed mitochondrial genes. **a**Repeated soft clustering of mitochondrial differentially expressed genes (DEGs) for the determination of an appropriate cluster number using *mfuzz* in R. **b** Principial component (PC) analysis plot of the six mitochondrial gene cluster centres that were segregated using *mfuzz* in R. **c** Representative genes involved in mitochondrial protein import from each mitochondrial cluster, indicative of a wide diversity of exercise-induced gene expression even within the same mitochondrial pathway. The membership score of each gene to its specific cluster is displayed in the title. **d** Heatmaps of mitochondrial cluster differential expression relative to the PRE time point compared to training-induced changes in protein abundance (PH vs BL comparison of Granata *et al*., no adjusted p-value threshold for inclusion)^26^. Row clustering determined by unsupervised hierarchical cluster analysis of the RNA-seq data.

**Extended Data Fig. 4:**
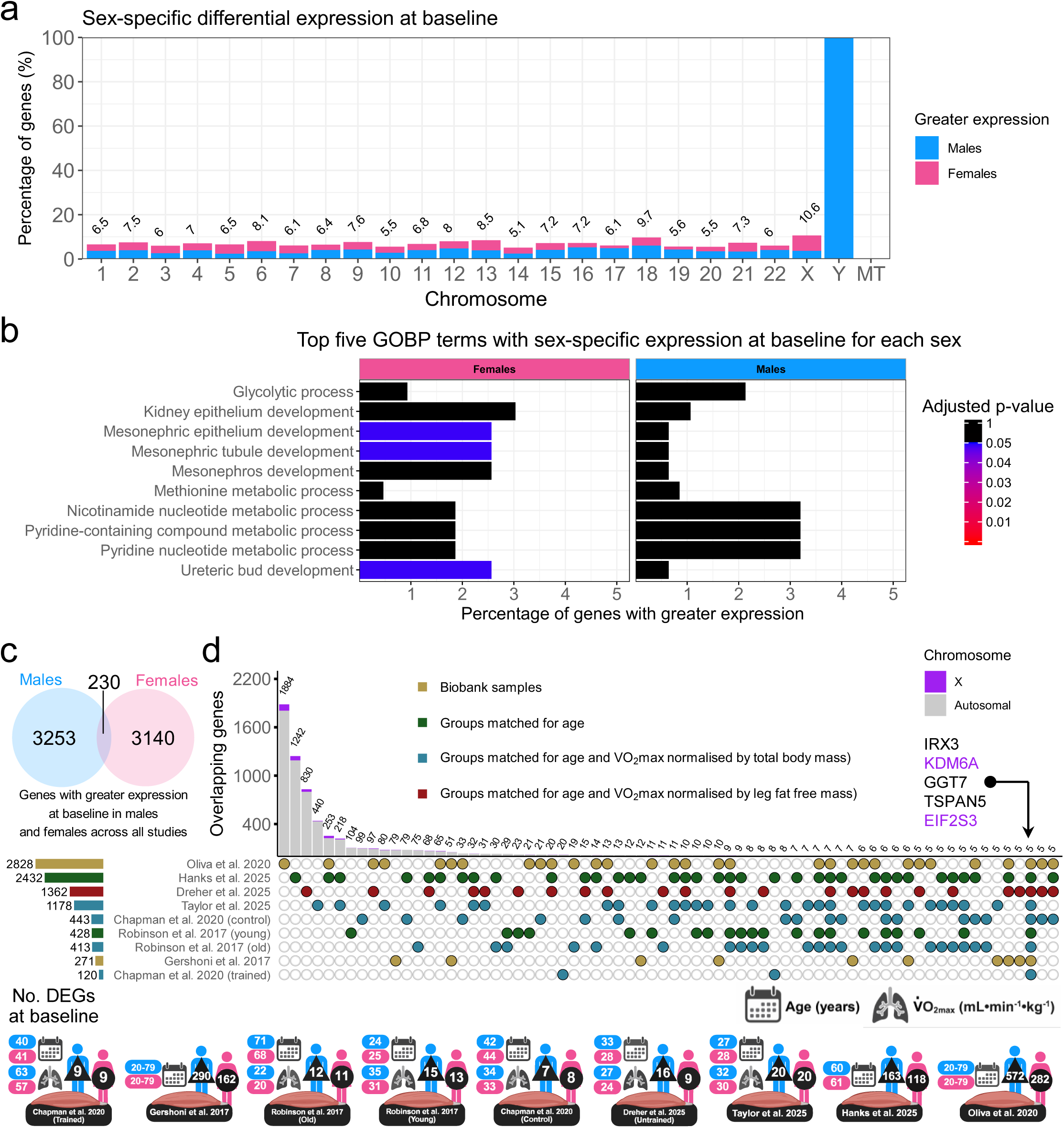
Effect of sex on baseline skeletal muscle gene expression. **a**Plot of chromosomal origin for genes with greater expression at baseline for each sex as a percentage of detected genes. MT: mitochondrial. **b** Gene ontology biological process (GOBP) enrichment for genes with sex-specific differential expression at baseline, significant enrichment determined with an adjusted p-value < 0.05 (Benjamini–Hochberg). **c** Venn diagram comparing genes identified as having greater baseline expression in males or females across all studies examining baseline sex-specific differences in skeletal muscle using RNA-seq. **d** Upset plot comparing differentially expressed genes (DEGs) detected within studies investigating baseline sex-specific expression in skeletal muscle using RNA-seq. Despite over 4,000 genes being identified as possessing sex-specific expression at baseline, only five genes were detected across all studies (excluding genes encoded on the Y chromosome). Minimum set size displayed is 5.

**Extended Data Fig. 5:**
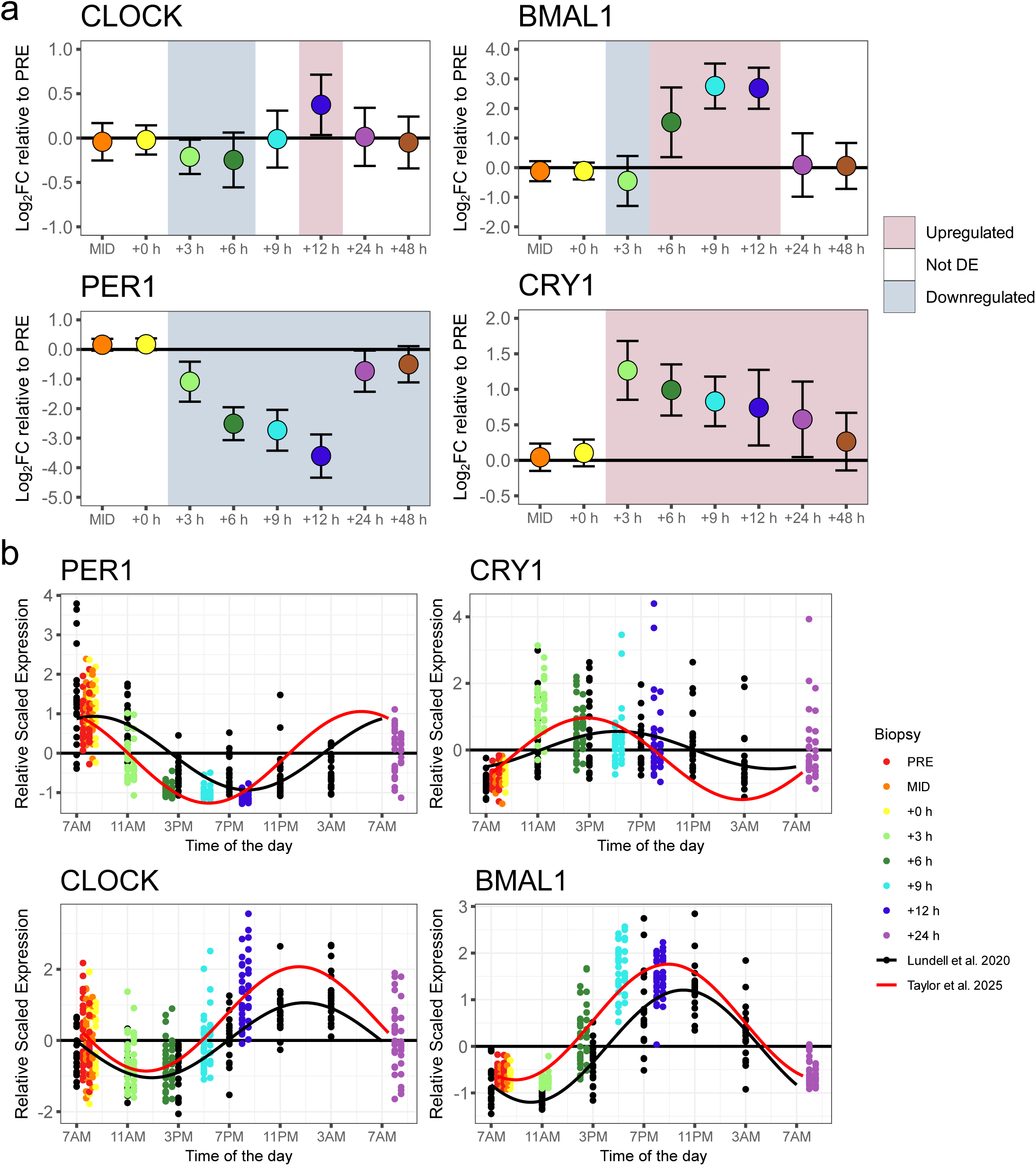
Effect of exercise on circadian gene expression in skeletal muscle. **a**Differential expression (determined by an adjusted p-value < 0.05 (Benjamini-Hochberg)) relative to PRE of circadian genes *CLOCK* and *BMAL1*, which form a heterodimer to regulate the expression of *PER1* and *CRY1*, which form a negative feedback loop to repress the expression of circadian genes. **b** Z-scored normalised gene expression for circadian genes *CLOCK*, *BMAL1*, *PER1*, and *CRY1* within this study compared to Lundell *et al.* 2020. Points are individual participants datapoints and lines represent a cosinor regression model fit assuming a periodicity of 24 hours.

**Extended Data Fig. 6:**
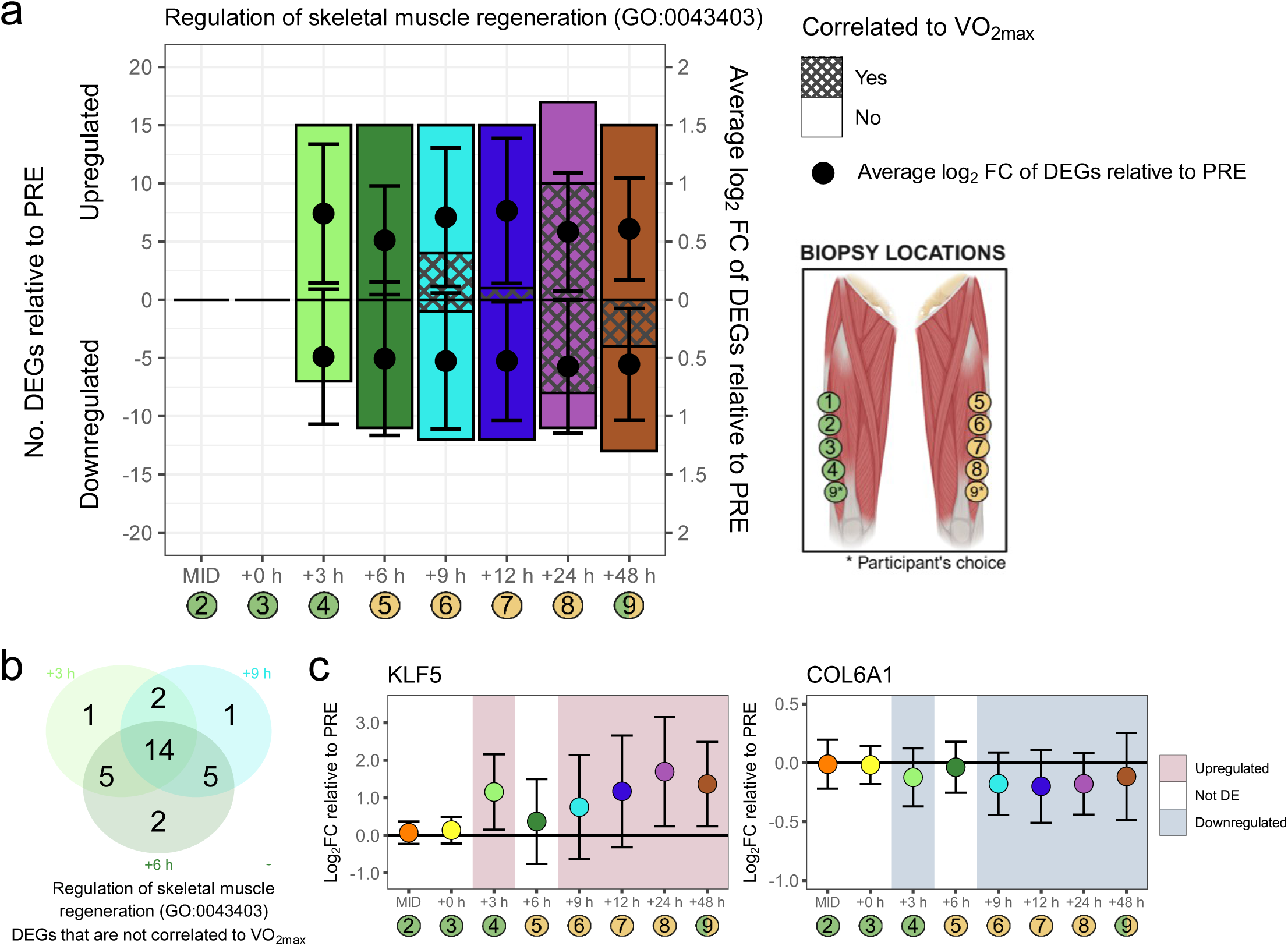
Effect of repeated biopsies on skeletal muscle gene expression. **a**The number of upregulated and downregulated differentially expressed genes (DEGs) relative to PRE at each subsequent time point within the GOBP term ‘regulation of skeletal muscle regeneration’ (GO:0043403), determined with an adjusted p-value < 0.05 (Benjamini-Hochberg). The number of DEGs with differential expression correlated to V̇O_2max_, and thus, induced by exercise and not the muscle biopsy, is also indicated. Additionally, the average log_2_ fold change (FC) of upregulated and downregulated DEGs is indicated. **b** Venn diagram of the overlap of DEGs within the GOBP term ‘regulation of skeletal muscle regeneration’ (GO:0043403) at +3 h, +6 h, and +9 h. The differential expression of genes at +3 h and +9 h, but not +6 h, potentially caused by the muscle biopsy. **c** Differential expression relative to PRE of *KLF5* and *COL6A1*, indicative of the upregulation of localised skeletal muscle repair in response to the skeletal muscle biopsies.

**Extended Data Fig. 7:**
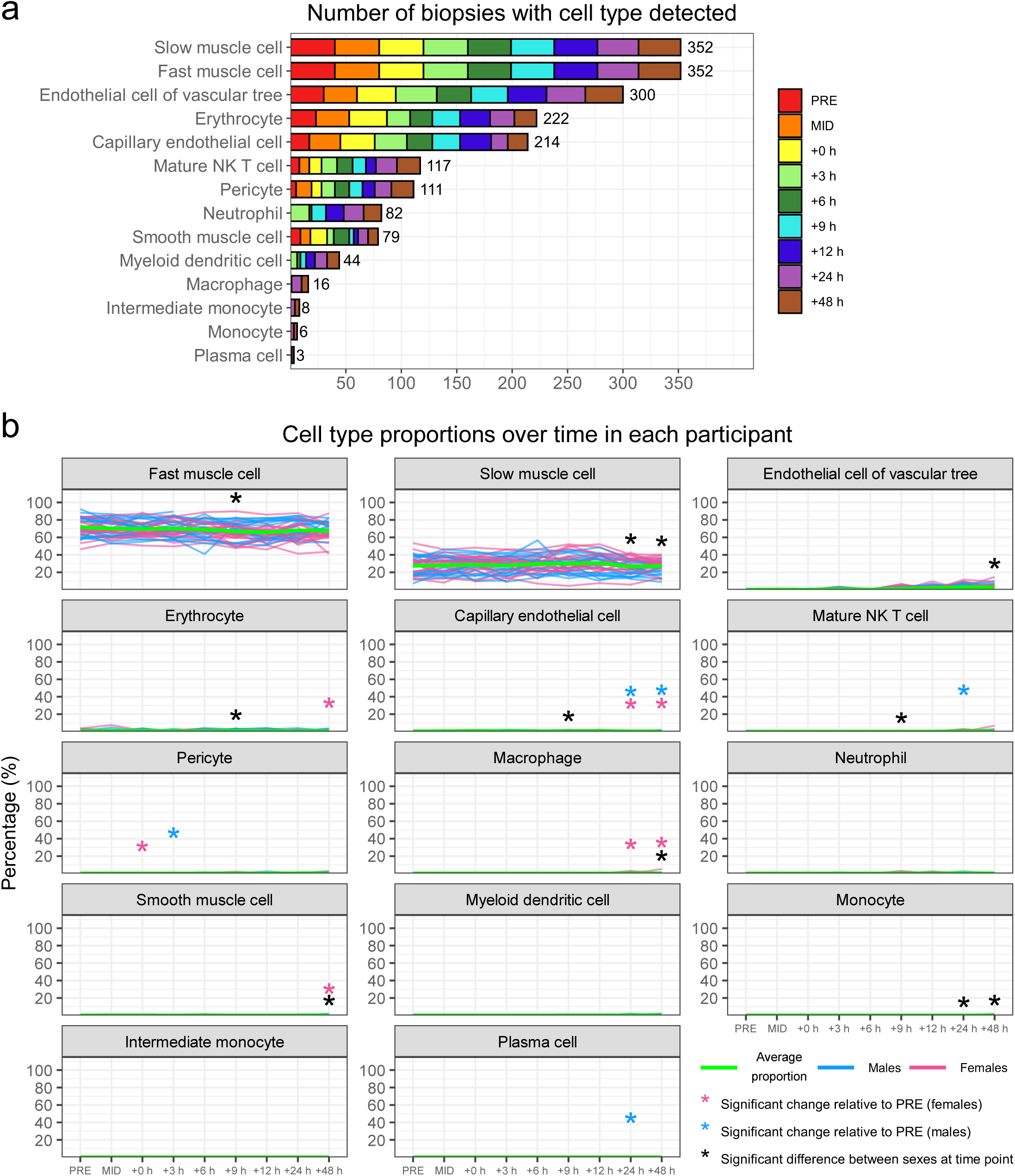
Sample cell type deconvolution. **a**The number of cell types identified at each time point through cell type deconvolution of normalised counts per million (CPM) data using the *MuSiC* package and the Tabula Sapiens single-cell RNA-seq atlas as a reference. **b** Percentage of bulk RNA-seq data attributed to each cell type across each time point in males and females. Significance determined by an adjusted p-value < 0.05 (Šidák correction).

**Extended Data Fig. 8:**
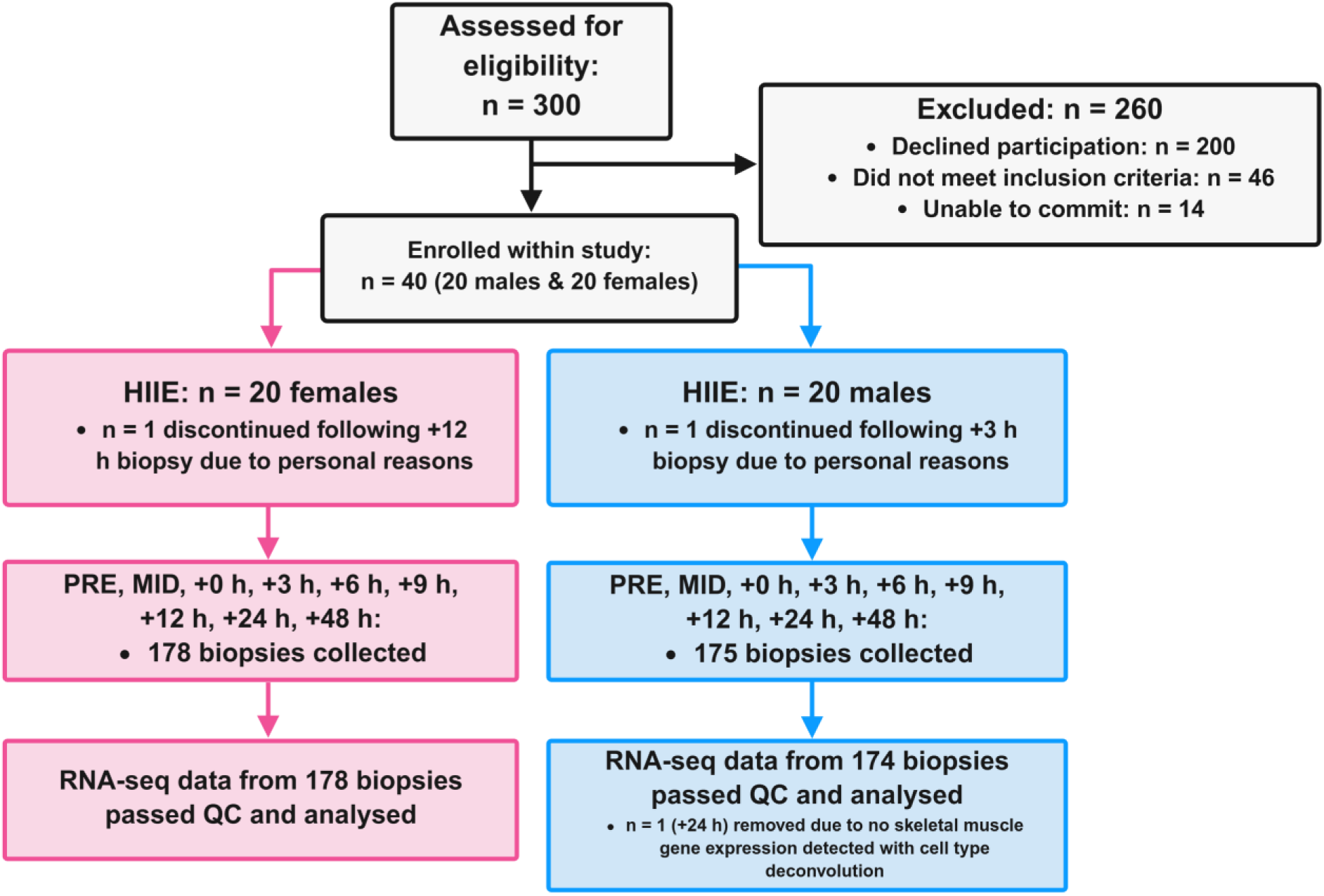
Participant recruitment consort flow chart. HIIE: high-intensity interval exercise.

## References

1 Hawley, J. A., Hargreaves, M., Joyner, M. J. & Zierath, J. R. Integrative biology of exercise. Cell 159, 738–749 (2014). 10.1016/j.cell.2014.10.029

2 Egan, B. & Sharples, A. P. Molecular responses to acute exercise and their relevance for adaptations in skeletal muscle to exercise training. Physiol Rev 103, 2057–2170 (2023). 10.1152/physrev.00054.2021

3 Reisman, E. G., Hawley, J. A. & Hoffman, N. J. Exercise-Regulated Mitochondrial and Nuclear Signalling Networks in Skeletal Muscle. Sports Med 54, 1097–1119 (2024). 10.1007/s40279-024-02007-2

4 Taylor, D. F. & Bishop, D. J. Transcription Factor Movement and Exercise-Induced Mitochondrial Biogenesis in Human Skeletal Muscle: Current Knowledge and Future Perspectives. Int J Mol Sci 23 (2022). 10.3390/ijms23031517

5 Bishop, D. J. & Hawley, J. A. Reassessing the relationship between mRNA levels and protein abundance in exercised skeletal muscles. Nat Rev Mol Cell Biol 23, 773–774 (2022). 10.1038/s41580-022-00541-3

6 Bishop, D. J. et al. Discordant skeletal muscle gene and protein responses to exercise. Trends Biochem Sci 48, 927–936 (2023). 10.1016/j.tibs.2023.08.005

7 Popov, D. V. et al. Intensity-dependent gene expression after aerobic exercise in endurance-trained skeletal muscle. Biol Sport 35, 277–289 (2018). 10.5114/biolsport.2018.77828

8 Popov, D. V. et al. Contractile activity-specific transcriptome response to acute endurance exercise and training in human skeletal muscle. Am J Physiol Endocrinol Metab 316, E605–E614 (2019). 10.1152/ajpendo.00449.2018

9 Dickinson, J. M. et al. Transcriptome response of human skeletal muscle to divergent exercise stimuli. J Appl Physiol (1985) 124, 1529–1540 (2018). 10.1152/japplphysiol.00014.2018

10 Rubenstein, A. B. et al. Skeletal muscle transcriptome response to a bout of endurance exercise in physically active and sedentary older adults. Am J Physiol Endocrinol Metab 322, E260–E277 (2022). 10.1152/ajpendo.00378.2021

11 Moore, T. M. et al. Conserved multi-tissue transcriptomic adaptations to exercise training in humans and mice. Cell Rep 42, 112499 (2023). 10.1016/j.celrep.2023.112499

12 Norrbom, J. M. et al. A HIF-1 signature dominates the attenuation in the human skeletal muscle transcriptional response to high-intensity interval training. J Appl Physiol (1985) 132, 1448–1459 (2022). 10.1152/japplphysiol.00310.2021

13 Reitzner, S. M. et al. Molecular profiling of high-level athlete skeletal muscle after acute endurance or resistance exercise - A systems biology approach. Mol Metab 79, 101857 (2024). 10.1016/j.molmet.2023.101857

14 Kuang, J. et al. Interpretation of exercise-induced changes in human skeletal muscle mRNA expression depends on the timing of the post-exercise biopsies. PeerJ 10, e12856 (2022). 10.7717/peerj.12856

15 Costello, J. T., Bieuzen, F. & Bleakley, C. M. Where are all the female participants in Sports and Exercise Medicine research? Eur J Sport Sci 14, 847–851 (2014). 10.1080/17461391.2014.911354

16 MacGregor, K., Ellefsen, S., Pillon, N. J., Hammarstrom, D. & Krook, A. Sex differences in skeletal muscle metabolism in exercise and type 2 diabetes mellitus. Nat Rev Endocrinol 21, 166–179 (2025). 10.1038/s41574-024-01058-9

17 Robinson, M. M. et al. Enhanced Protein Translation Underlies Improved Metabolic and Physical Adaptations to Different Exercise Training Modes in Young and Old Humans. Cell Metab 25, 581–592 (2017). 10.1016/j.cmet.2017.02.009

18 Gershoni, M. & Pietrokovski, S. The landscape of sex-differential transcriptome and its consequent selection in human adults. BMC Biol 15, 7 (2017). 10.1186/s12915-017-0352-z

19 Chapman, M. A. et al. Skeletal Muscle Transcriptomic Comparison between Long-Term Trained and Untrained Men and Women. Cell Rep 31, 107808 (2020). 10.1016/j.celrep.2020.107808

20 Oliva, M. et al. The impact of sex on gene expression across human tissues. Science 369 (2020). 10.1126/science.aba3066

21 Dreher, S. I. et al. Sex differences in resting skeletal muscle and the acute and long-term response to endurance exercise in individuals with overweight and obesity. Mol Metab 98, 102185 (2025). 10.1016/j.molmet.2025.102185

22 Hanks, S. C. et al. Extensive differential gene expression and regulation by sex in human skeletal muscle. Cell Genom, 100915 (2025). 10.1016/j.xgen.2025.100915

23 Pillon, N. J. et al. Transcriptomic profiling of skeletal muscle adaptations to exercise and inactivity. Nat Commun 11, 470 (2020). 10.1038/s41467-019-13869-w

24 Amar, D. et al. Time trajectories in the transcriptomic response to exercise - a meta- analysis. Nat Commun 12, 3471 (2021). 10.1038/s41467-021-23579-x

25 Botella, J. et al. Sprint interval exercise disrupts mitochondrial ultrastructure driving a unique mitochondrial stress response and remodelling in humans. bioRxiv (2024). 10.1101/2024.12.19.629456

26 Granata, C., Oliveira, R. S. F., Little, J. P. & Bishop, D. J. Forty high-intensity interval training sessions blunt exercise-induced changes in the nuclear protein content of PGC-1alpha and p53 in human skeletal muscle. Am J Physiol Endocrinol Metab 318, E224–E236 (2020). 10.1152/ajpendo.00233.2019

27 Granata, C. et al. High-intensity training induces non-stoichiometric changes in the mitochondrial proteome of human skeletal muscle without reorganisation of respiratory chain content. Nat Commun 12, 7056 (2021). 10.1038/s41467-021-27153-3

28 Furrer, R. et al. Molecular control of endurance training adaptation in male mouse skeletal muscle. Nat Metab 5, 2020–2035 (2023). 10.1038/s42255-023-00891-y

29 Bishop, D. J., Lee, M. J. & Picard, M. Exercise as Mitochondrial Medicine: How Does the Exercise Prescription Affect Mitochondrial Adaptations to Training? Annu Rev Physiol 87, 107–129 (2025). 10.1146/annurev-physiol-022724-104836

30 Williams, R. S. N., P. D. Regulation of gene expression in skeletal muscle by contractile activity., 1124–1150 (American Physiological Society, 1996).

31 Hood, D. A. Invited Review: contractile activity-induced mitochondrial biogenesis in skeletal muscle. J Appl Physiol (1985) 90, 1137–1157 (2001). 10.1152/jappl.2001.90.3.1137

32 Perry, C. G. et al. Repeated transient mRNA bursts precede increases in transcriptional and mitochondrial proteins during training in human skeletal muscle. J Physiol 588, 4795–4810 (2010). 10.1113/jphysiol.2010.199448

33 Leick, L. et al. Endurance exercise induces mRNA expression of oxidative enzymes in human skeletal muscle late in recovery. Scand J Med Sci Sports 20, 593–599 (2010). 10.1111/j.1600-0838.2009.00988.x

34 Maleki, F., Ovens, K., McQuillan, I. & Kusalik, A. J. Size matters: how sample size affects the reproducibility and specificity of gene set analysis. Hum Genomics 13, 42 (2019). 10.1186/s40246-019-0226-2

35 Emanuelsson, E. B. et al. Remodeling of the human skeletal muscle proteome found after long-term endurance training but not after strength training. iScience 27, 108638 (2024). 10.1016/j.isci.2023.108638

36 Nie, M., Liu, Q. & Yan, C. Skeletal Muscle Transcriptomic Comparison Between Men and Women in Response to Acute Sprint Exercise. Front Genet 13, 860815 (2022). 10.3389/fgene.2022.860815

37 Landen, S. et al. Sex differences in muscle protein expression and DNA methylation in response to exercise training. Biol Sex Differ 14, 56 (2023). 10.1186/s13293-023-00539-2

38 Baldwin, J., Snow, R. J. & Febbraio, M. A. Effect of training status and relative exercise intensity on physiological responses in men. Med Sci Sports Exerc 32, 1648–1654 (2000). 10.1097/00005768-200009000-00020

39 Benitez-Munoz, J. A. et al. Greater Relative First and Second Lactate Thresholds in Females Compared With Males: Consideration for Exercise Prescription. Int J Sports Physiol Perform, 1–7 (2024). 10.1123/ijspp.2024-0079

40 Martin, R. A., Viggars, M. R. & Esser, K. A. Metabolism and exercise: the skeletal muscle clock takes centre stage. Nat Rev Endocrinol 19, 272–284 (2023). 10.1038/s41574-023-00805-8

41 Lundell, L. S. et al. Time-restricted feeding alters lipid and amino acid metabolite rhythmicity without perturbing clock gene expression. Nat Commun 11, 4643 (2020). 10.1038/s41467-020-18412-w

42 Edman, S. et al. The 24-Hour Time Course of Integrated Molecular Responses to Resistance Exercise in Human Skeletal Muscle Implicates MYC as a Hypertrophic Regulator That is Sufficient for Growth. bioRxiv (2024). 10.1101/2024.03.26.586857

43 Van Thienen, R., D’Hulst, G., Deldicque, L. & Hespel, P. Biochemical artifacts in experiments involving repeated biopsies in the same muscle. Physiol Rep 2, e00286 (2014). 10.14814/phy2.286

44 Friedmann-Bette, B. et al. Similar changes of gene expression in human skeletal muscle after resistance exercise and multiple fine needle biopsies. J Appl Physiol (1985) 112, 289–295 (2012). 10.1152/japplphysiol.00959.2011

45 Vissing, K., Andersen, J. L. & Schjerling, P. Are exercise-induced genes induced by exercise? FASEB J 19, 94–96 (2005). 10.1096/fj.04-2084fje

46 Guerra, B. et al. Repeated muscle biopsies through a single skin incision do not elicit muscle signaling, but IL-6 mRNA and STAT3 phosphorylation increase in injured muscle. J Appl Physiol (1985) 110, 1708–1715 (2011). 10.1152/japplphysiol.00091.2011

47 Van de Casteele, F. et al. Does one biopsy cut it? Revisiting human muscle fiber type composition variability using repeated biopsies in the vastus lateralis and gastrocnemius medialis. J Appl Physiol (1985) 137, 1341–1353 (2024). 10.1152/japplphysiol.00394.2024

48 D’Souza, A. C. et al. Menstrual cycle hormones and oral contraceptives: a multimethod systems physiology-based review of their impact on key aspects of female physiology. J Appl Physiol (1985) 135, 1284–1299 (2023). 10.1152/japplphysiol.00346.2023

49 Reisman, E. G. et al. Fibre-specific mitochondrial protein abundance is linked to resting and post-training mitochondrial content in the muscle of men. Nat Commun 15, 7677 (2024). 10.1038/s41467-024-50632-2

50 Jamnick, N. A., Botella, J., Pyne, D. B. & Bishop, D. J. Manipulating graded exercise test variables affects the validity of the lactate threshold and V̇O2peak. PLoS One 13, e0199794 (2018). 10.1371/journal.pone.0199794

